# Coordination of gene expression with cell size enables *Escherichia coli* to efficiently maintain motility across conditions

**DOI:** 10.1101/2021.05.12.443892

**Authors:** Tomoya Honda, Jonas Cremer, Leonardo Mancini, Zhongge Zhang, Teuta Pilizota, Terence Hwa

**Author notes:** These authors contributed equally to this work (Tomoya Honda, Jonas Cremer).

## Abstract

To swim and navigate, motile bacteria synthesize a complex motility machinery involving flagella, motors, and a sensory system. A myriad of studies has elucidated the molecular processes involved, but less is known about the coordination of motility expression with cellular physiology: In *Escherichia coli*, motility genes are strongly upregulated in nutrient-poor conditions compared to nutrient-replete conditions; yet a quantitative link to cellular motility has not been developed. Here, we systematically investigate gene expression, swimming behavior, and cell growth across a broad spectrum of exponential growth condition. We establish that *E. coli* up-regulates the expression of motility genes at slow growth to compensate for reduction in cell size, such that the number of flagella per cell is maintained across conditions. The observed 4-5 flagella per cell is the minimum number needed to keep the majority of cells motile. This simple regulatory objective allows *E. coli* cells to remain motile across a broad range of growth conditions while keeping the biosynthetic and energetic demands to establish and drive the motility machinery at the minimum needed. Given the strong reduction in flagella synthesis resulting from cell size increases at fast growth, our findings also provide a novel physiological perspective on bacterial cell size control: A larger cell-size at fast growth is an efficient strategy to increase the allocation of cellular resources to the synthesis of those proteins required for fast growth, while maintaining processes such as motility which are only needed on a per-cell basis.

---

To thrive in different environments, bacteria must efficiently allocate their limited resources towards different cellular processes in accordance to what is most needed for their growth and survival (1). Flagella driven motility is one of the most distinct processes of bacterial life which provides cells with novel ways to respond to the conditions they encounter (2). The active movement towards more favorable conditions and away from detrimental ones has been studied in detail on the molecular level (3–5) and can give rise to strong fitness advantages (6–9). But flagella driven motility is also a resource demanding process. For growing *E.coli* cells, the synthesis of the motility proteins alone ties up a substantial portion of the protein synthesis resources (10, 11), and the assembly and rotation of flagella also demand energy (12–14). Accordingly, motility expression constitutes a burden on cell growth, such that cells with attenuated motility can grow up to 20 % faster and reach about 10 % higher biomass yields (15–17), a strong difference readily affecting the outcome of (laboratory) evolution (18–21). Given this burden, the expression of motility is expected to be highly controlled, coordinated with other cellular processes and demands.

Notably, the expression of motility genes varies strongly with the nutrient conditions cells encounter and more resources are allocated to motility expression in nutrient poor than in replete conditions (22–26). These observations have been taken as support for the idea that motility is a response expressed to search for alternative nutrient sources when local nutrient sources are depleted (22, 25, 26). However, swimming speeds observed during balanced growth do not vary much with the growth rate or the carbon source provided (9). Furthermore, bacterial population exhibit a chemotaxis-driven range expansion (6, 8, 27–29) with expansion speed which is markedly faster in nutrients providing faster growth (9). These latter observations suggest that motility is a phenotype broadly expressed by growing cells, rather than being merely being a foraging response to starvation. But then why are motility genes expressed more in poor growth conditions and how does their degree of expression quantitatively affect swimming? To resolve this puzzle, we systematically investigated the link between gene expression and swimming in different balanced growth conditions. We found that *E. coli* cells maintain motility by regulating gene expression in coordination with cell size; upregulation of motility genes at slower growth is a necessary compensation to adjust for growth-related changes in cell size such that the number of flagella per cell remains constant. This simple regulatory objective provides an example of how cells maintain function while keeping resource demands minimal. Our findings also provide a new perspective on the relation between cell size control and proteome resource allocation, giving a physiological rationale for the ubiquitously observed positive relation between cell growth and cell size.

To study the relation between swimming behavior and motility gene expression, we first examined gene expression during balanced growth across a broad range of growth conditions, using a physiologically well-characterized strain (WT strain *E. coli* K-12 HE204, **SI Text 1.1**). Motility genes are hierarchically regulated and have been assigned into three different classes with the master regulator *flhDC* being the class-I genes as illustrated in **Fig. S1A** (30–32). We first studied the expression of *fliA*, a class-II gene which is expressed coordinately with other class-II genes (including those encoding flagella hook-basal body components,**Fig. S1BC**) and encodes the sigma factor *σ_F_* required for the expression of flagella components (class-III genes). Using a LacZ reporter, we quantified the expression level of the *fliA* promoter (in unit of LacZ activity per biomass; see **SI Text 1.4**) during balanced growth, with a range of growth rates obtained by supplementing minimal medium with different carbon sources or rich media components (detailed growth conditions described in **SI Text 1.2**). Consistent with previous reports (22, 24–26), *fliA* expression was higher at slower growth rates (**Fig. 1A**, circles): Expression levels change approximately exponentially with growth rate (dashed line), with a ~4.4 fold increase when growth rates change from fast (1.60 1/*h* for rich defined medium with glucose) to slow (0.28 1/*h* for aspartate). In contrast, a constitutively active promoter, reported by P*tet-lacZ* expression, exhibits only a ~1.3 fold change **(Fig. 1A**, diamond). The coordinated expression of *fliA* and other motility genes provides a substantial growth-rate dependent burden for the cell: a deletion of the master regulator *flhD*, resulting in the complete suppression of motility gene expression, increased growth rate by up to 18% compared to the WT strain, with larger increases realized in slower growth conditions where motility expression in the WT is higher (**Fig. S1D**).

**Fig. 1:**
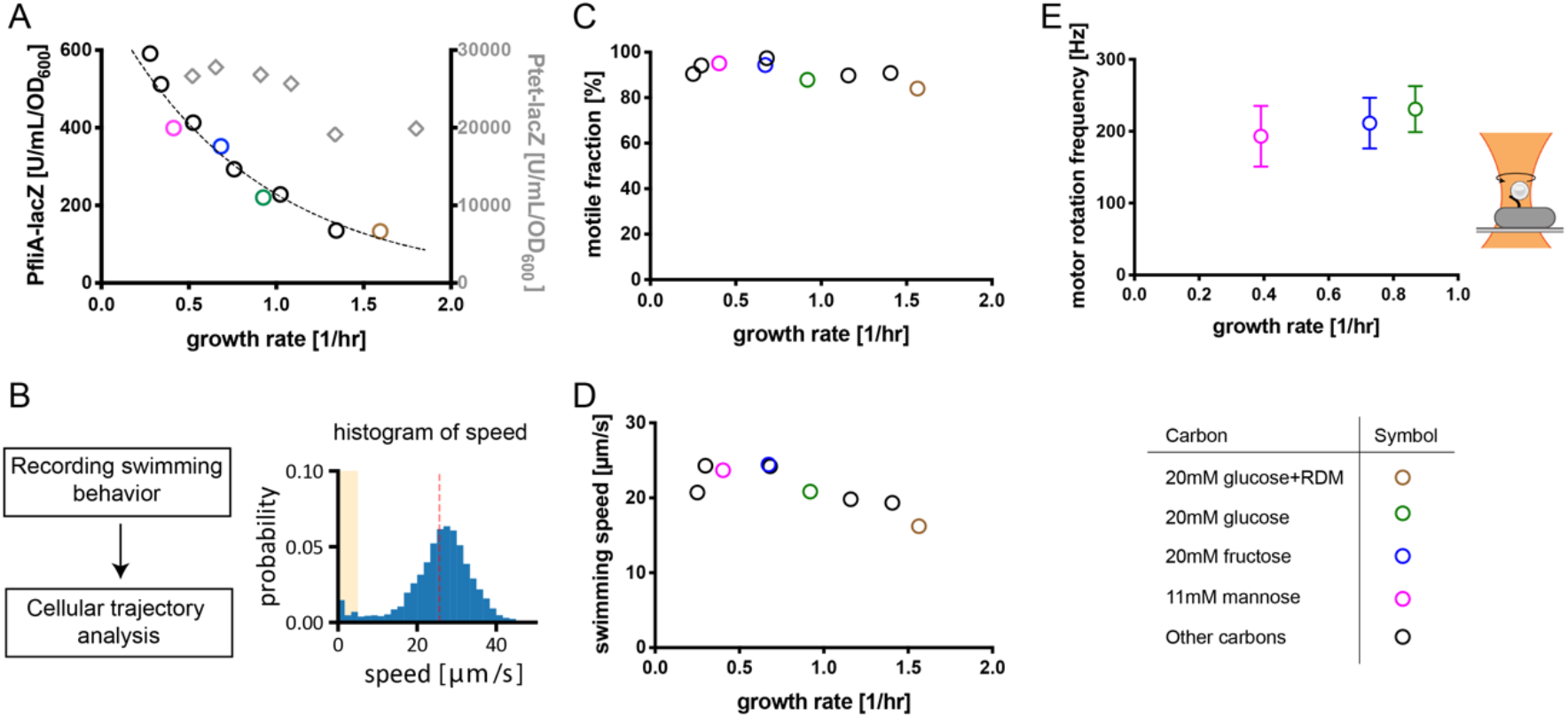
Motility gene expression and swimming behavior during balanced growth in different growth media. **(A)** Expression level of a reporter of the *fliA* promoter (a class-II gene; Fig. S1A) for growth on different carbon sources (circles). The reporter expression changes strongly with growth rates. Dashed line indicates exponential fit, 734 · *e^−k·λ^* with rate *k* = 1.17 1/*h*. In contrast, reporter expression driven by a constitutively active promoter, P*tet-lacZ* (strain NQ122), changes weakly (diamonds). **(B)** Quantification of swimming behavior: Using a phase contrast microscopy, cells were tracked and swimming speeds were analyzed (details in **SI Text 1.3**). Cell-to-cell variation of swimming speeds for growth on fructose as example. The red line indicates the mean swimming speed, and the yellow background indicates the range defined as non-motile (swimming speeds *v_i_* < 5 μ*m/s*). Additional conditions and reproducibility are shown in **Fig. S2**. **(C, D)** Motile fractions and average swimming speeds for different growth rates, which show minor variations with growth-rate. (**E)** Motor rotation frequency for different growth rates. Rotation frequencies of beads attached to filament stub were measured using back-focal plane interferometry and a strain with the filament gene modified to readily stick to polystyrene beads (sticky-*fliC*) (49, 50), see cartoon and **SI Text 1.6**. Data in rich media were not collected because rapid cell divisions prevented the motor observation for sufficient periods. Error bars indicate s.d. observed for the probed population. Four reference conditions are highlighted by colors as indicated in the legend. Strains HE207, HE206 and HE608 were used in (A), (C,D) and (E), respectively.

To understand the consequences of this costly expression on swimming, we next characterized the swimming behavior across growth conditions. Extending a previous approach combining phase contrast microscopy and tracking (9), we quantified the movement of hundreds of cells and analyzed the distributions of observed swimming speeds {*v_i_*}during run events (see **Fig. 1B, Fig. S2** and **SI Text 1.3** for methods). We then extracted the average swimming speed and the fraction of *motile cells* with swimming velocities *v_i_* > 5 *μm/s*. Notably, despite the ~4.4 fold change of gene expression (**Fig. 1A**), swimming characteristics varied only weakly: the fraction of swimming cells (*α_m_*) remained close to 90 % for all growth conditions (**Fig. 1C**), and the average swimming speed, 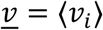, changed only ~1.3 fold from fast (rich defined medium with glucose) to slow growth (aspartate) (**Fig. 1D**).

One possible explanation for this combined observation of minor changes in motile behavior and the large changes in expression of motility genes would be an adjustment to a possible decrease in flagella motor activity at slow growth: The *E. coli* flagella motor is driven by the proton motive force (PMF) and the motor rotation frequency is proportional to the PMF (12, 13). Given that the PMF is a result of the metabolic state which might change with growth condition, the cell might compensate for slower rotation in poor growth conditions by increasing the expression level of motility genes. To probe this possibility, we measured the motor activity by tracking the rotation of beads attached to flagella filaments (33, 34). However, the rotation frequency is found to be almost independent of growth (**Fig. 1E**; a drop of 13 % from growth rate 0.87 1/*h* to 0.39 1/*h*).

Why then are motility genes expressed more in slow growth conditions? To investigate this question further, we next performed experiments with a synthetic construct which allows for the smooth titration of motility gene expression in a given growth condition, so that we can separately assess the effect of changing motility expression and growth. We replaced the native promoter of the master regulator *flhDC* by the P*tet* promoter, enabling an inducer-dependent control. Additionally, the construct also carries the above-mentioned P*fliA-lacZ* as a reporter for a class-II gene expression (see **Fig. 2A** and **SI Text 1.1.3** for cartoon and details). We first grew this strain in fructose minimal medium with different concentrations of the inducer chlortetracycline (cTc). P*fliA-lacZ* expression decreased smoothly from wild-type levels towards zero when reducing the inducer concentration in the media (**Fig. 2A** blue points). Decreasing the inducer concentration similarly shifted the distribution of swimming speeds (**Fig. 2B)**. with falling average swimming speeds and motile fractions (**Fig. 2CD,** blue points). Similar results were obtained by growing cells in other carbon sources that provide faster and slower growth rates (**Fig. 2,** glucose and mannose as green and magenta points). Overall, these results show that motility gene expression has a strong influence on cellular swimming behaviors at each growth condition, as can also be seen by directly plotting swimming speed and motility fraction against *fliA* expression (**Fig. S3**).

**Fig. 2:**
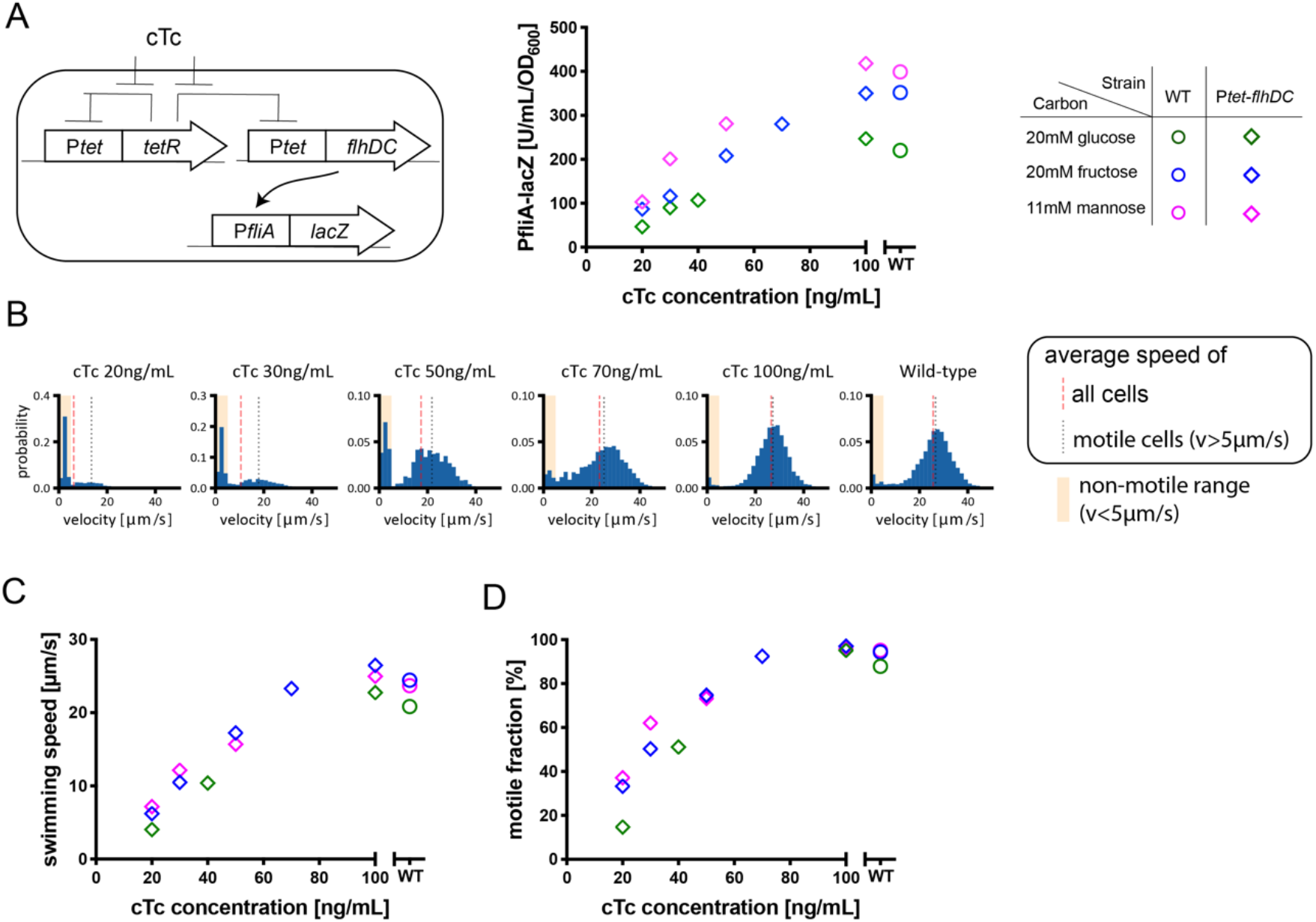
Gene expression and swimming behavior when titrating a motility master regulator. **(A)** Titratable *flhDC* construct with P*fliA*-*lacZ* reporter to quantify *fliA* expression. Titration control is achieved via the P*tet* system and cTc as inducer. P*fliA-lacZ* expression was measured by varying inducer concentration. (**B**) Swimming speed distributions when cells are grown on fructose with different inducer levels. (**C** & **D**) Changes in average swimming speed and the motile fraction of cells (swimming speed > 5 μm/s) in the population. WT are shown as circles in A, C, D for comparison. Strains HE641 and HE170 were used in (A) and (B,C,D), respectively (both strains are identical except carrying different *lacZ* reporters: **Table S1**).

To further study the role of growth-rate dependent regulation of motility genes on swimming behavior, we next compared how the swimming phenotypes change at given levels of motility gene expression (independent of the growth rate) using the titratable construct and selected inducer levels, as indicated by the dotted and dashed lines in **Fig. 3A**. Comparing the swimming behavior at these fixed expression levels, we found a gradual reduction of swimming speed as the growth rate slows down (**Fig. 3BC**). The decreases in swimming speed at fixed expression levels can be largely accounted for by reduced fraction of motile cells (**Fig. 3D**), while the average swimming speed of those motile cells did not change significantly (**Fig. 3B**, grey vertical lines). Together, these observations establish that the upregulation of motility genes at slower growth is necessary to keep the population motile but not to increase the swimming speed of the motile cells.

**Fig. 3:**
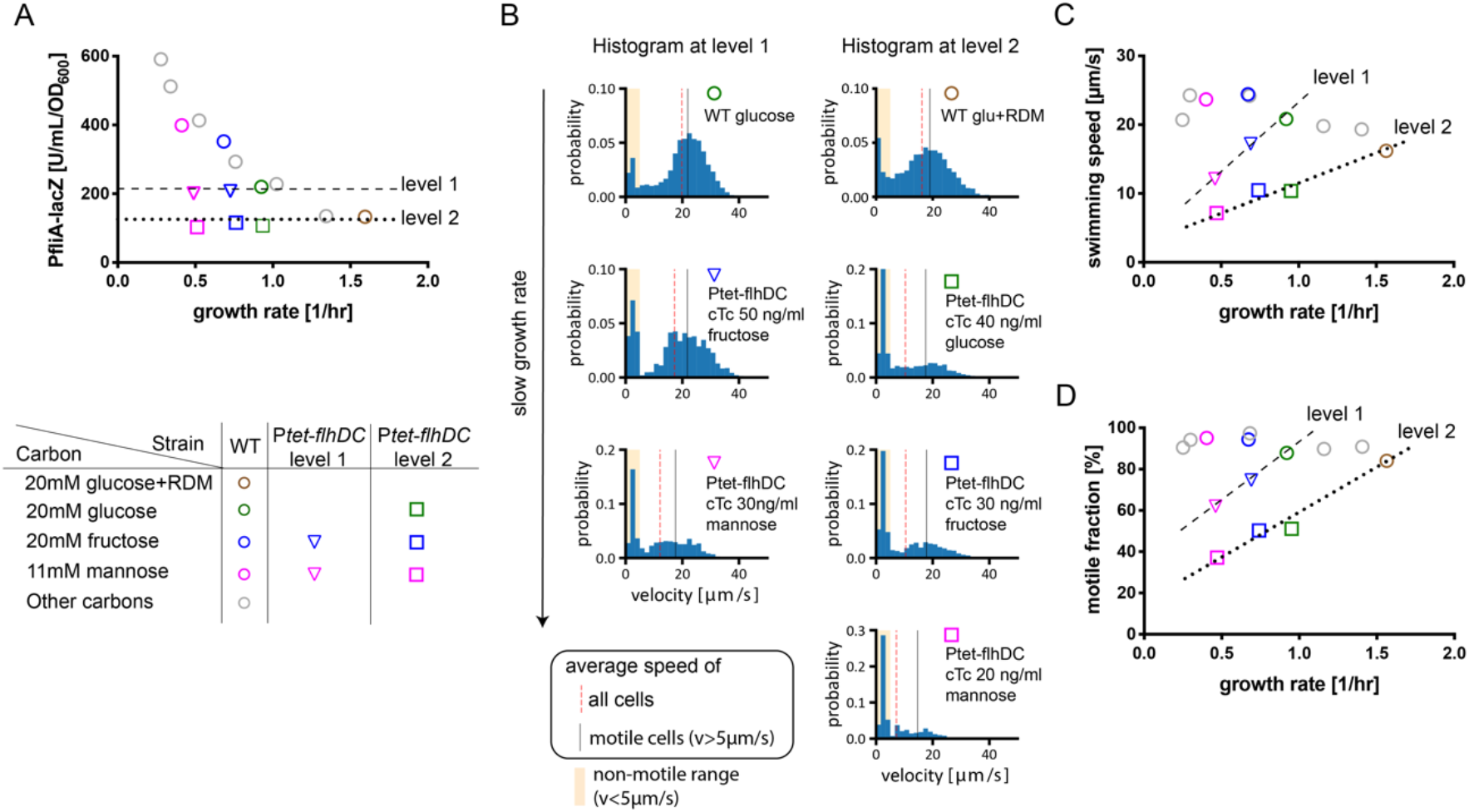
Swimming behavior across growth conditions at fixed motility expression. (**A**) P*fliA-lacZ* expression against growth rate when *flhDC* levels are titrated. Colors indicate three selected carbon sources (see legend). Two different expression levels, close to the levels observed for WT growth in glucose (level 1) and glucose+RDM (level 2) were chosen to study swimming behavior at fixed expression levels independent of growth rate. (**B**) Distributions of swimming speed for the two expression levels (left and right columns). The fraction of non-motile cells increases for decreasing growth rate (bars in yellow regions), while the average swimming speed of motile cells (grey lines) barely changes. **(C, D)**. The average swimming speed (D) and motile fraction (E) drop for the two fixed expression levels as growth rates decrease (dashed and dotted lines), while WT cells (circles) exhibit minor changes. Strain HE641 and HE170 were used in (A) and (B,C,D), respectively.

To better understand the regulation of motility genes and its connection to swimming behavior, we next considered the abundance of motility gene products per cell: Gene expression levels, as those determined via a LacZ reporter, are typically quantified per biomass (*e.g*., the commonly used “Miller Unit” (35) quantifies LacZ activity per OD, *U*/ *ml*/OD_600_, with OD_600_ having a constant relation with biomass across growth condition (36); see **SI Text 1.4 & SI Text 2)**. Since biomass itself is proportional to cell volume due to the constancy of biomass density (37, 11), the measurements with the class-II gene reporter P*fliA-lacZ* reflect the *concentration* of class-II gene products (flagella hook & basal body; **Fig. S1, Fig. 4A**, top row). As previously discussed, this concentration is higher when cells grow slower (**Fig. 1A**). However, bacterial cells also have different cell sizes at different growth rates. In fact, the biomass per cell exhibits an approximate exponential dependence on the growth rate (**Fig. 4B**), known as the Schaechter-Maaloe-Kjeldgaard relation (38–40). Accordingly, the *abundance of class-II gene products per cell* is expected to exhibit less change with growth rate than what is observed for the *concentration* (**Fig. 4A**, bottom row). Confirming this idea, the P*fliA-lacZ* expression per cell (unit: *U* /*cell*), taken as the product of expression per biomass (unit: *U / ml/OD*_600_) and the biomass per cell (unit: *ml* · *OD*_600_/*cell*), is nearly independent of growth rate (**Fig. 4C**, filled red points). Remarkably, the exponential relations observed for cell size (**Fig. 4B**, dashed line) and the expression level per biomass (**Fig. 4C**, dashed black line) show similar absolute rates (1.18 *h*^−1^ and 1.17 *h*^−1^), leading to the abundance per cell being independent of growth rate (**Fig. 4C**, dotted red line).

**Fig. 4:**
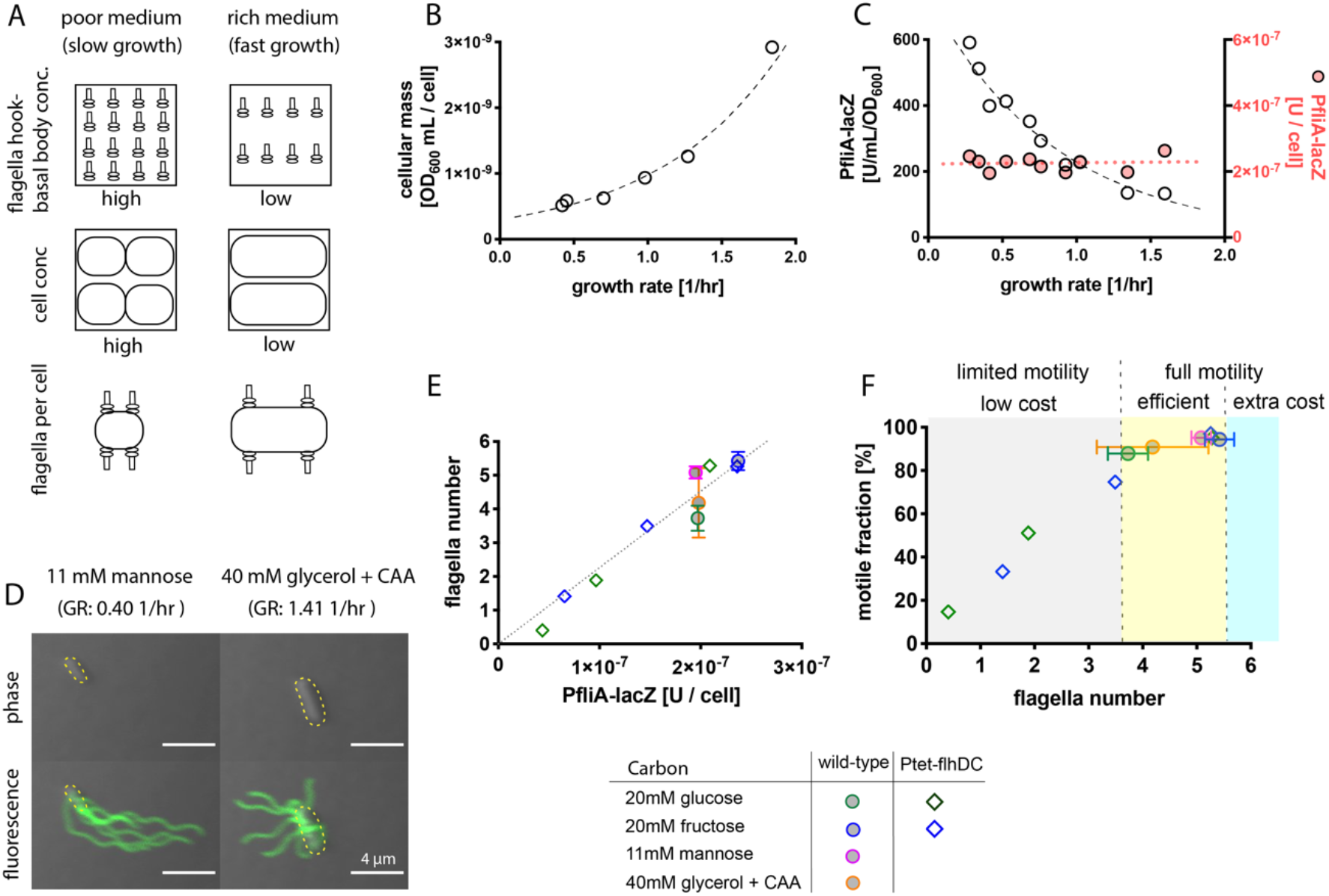
Expression levels change with cell size such that cells remain motile across growth conditions. **(A)** The concentration of motility gene products per unit biomass is higher for growth in poor condition (top), but cells are also smaller when growing slower (middle). Consequently, the abundance of gene products expressed per cell, like the number of flagella, would show less variation across growth conditions (bottom). (B) The biomass of cells, determined via optical density measurements and cell counting (CFU in culture), increases exponentially with growth rate. Line indicates exponential fit, ~*e^k·λ^*, with growth rate *λ* and parameter *k* = 1.18 1/*h*. (C) Expression of the class-II gene *fliA* per biomass (open circles) and per cell (red circles) based on the P*fliA-lacZ* measurements (strain HE207). Dashed line shows an exponential fit for the expression per biomass, ~*e^k·λ^* with rate *k* = −1.17 1/*h*. A red dotted line indicates the product of the two exponential relations in cell size (B) and P*fliA-lacZ* per biomass. (D) Images of cells with stained flagella filaments in a rich and poor condition (slow and fast growth, left and right column). While size differs (phase contrast, top row), a similar number of flagella filaments is observed (fluorescently labeled filaments, bottom row). Full distribution across the population shown in Fig. S4. Yellow lines indicate cell perimeter. (E) Relation between flagella filament number and P*fliA-lacZ* expression per cell for the native (filled circles) and titratable (diamonds) regulation of *flhDC* expression. Line indicates a linear fit with slope 2.26 · 10^7^. (F) Motile fraction and flagella number for wild-type (circles) and titratable *flhDC* strain (diamonds). WT cells are mostly motile and adjust their expression level per cell to ensure motility (yellow region) while preventing more expression than needed (blue region). Strains HE206 and HE170 used to obtain swimming data. Strains HE207 and HE641 used to obtain P*fliA-lacZ* expression data. Flagella filament numbers were quantified using strains HE582 and HE571 harboring a modified S219C *fliC* sequence. Cell size data in (B) from Basan *et al* (36).

The above analysis suggests that cells maintain their *number* of flagella across growth conditions and that the large change of gene expression with growth rate is necessary to keep this number constant as the cell size changes. To more directly confirm this idea, we counted the number of flagella filaments attached to the cells using a staining assay (**Fig. S4** and **SI Text 1.5**). We confirmed that the average number of flagella filaments per WT cell remains within a narrow range across growth conditions (4-5, within the measurement error), see **Fig. S4D**. As an example, two cells of different sizes but similar flagella numbers are shown in **Fig. 4D**. Looking at the distribution of filament numbers across the population, we see that very few cells possess only one or zero filaments (**Fig. S4B**), consistent with a high fraction of motile cells (**Fig. 1C**). In contrast, the average number of filaments varied strongly for the titratable *flhDC* strain as the provided inducer concentration was varied (**Fig. S5**). Particularly, the fraction of cells with zero or one filament clearly increased at lower inducer concentrations (**Fig. S5AB**) which coincides with the increase in the fraction of non-motile cells at lower inducer concentrations (**Fig. 2D**). We further confirmed that the class-II gene reporter expression reflects the change of filament number (**Fig. 4E**): reducing P*fliA-lacZ* level by titrating *flhDC* expression led to a linear drop of the average number of filaments in different growth conditions (**Fig. 4E**, open symbols). In contrast, the WT strain exhibited little variation in either the filament number or gene expression per cell (**Fig. 4E**, filled circles). In combination, our findings reveal that the regulated adjustment of motility gene expression in different growth conditions compensates for the changes in cell size seen in these conditions, such that a similar number of flagella is maintained for each cell across conditions.

To see how efficiently the motility genes are regulated, consider the relation between the number of flagella per cell, and the fraction of motile cells (**Fig. 4F**): When the gene expression is low such that there are < 4 flagella/cell (*flhDC* titration with low inducer levels, diamonds), the motile fraction is proportional to the flagella number (**Fig. 4F**, grey region: limited motility). In contrast, when expression levels reach close to those of WT such that there are > 4 flagella/cell, almost all cells are motile (**Fig. 4F**, circle points and yellow region: full motility). An even higher expression level per cell would only increase the costs to express extra flagella and is not observed (**Fig. 4F**, blue region: non-efficient expression). *E. coli* thus appears to regulate its motility expression levels such that the associated resource demands to synthesize and rotate flagella are at the minimum necessary to keep most cells motile. While the requirement for ~4 flagella per cell ensures most cells to be motile (yellow region in **Fig. 4F**), this number is also close to what is minimally required to allow uninterrupted motility when cells half the number of their flagella during cell division.

In this study, we analyzed the regulation of motility genes by *E. coli* for different balanced growth conditions and found that the fold-change in gene expression per biomass compensates for the variation in cell size, resulting in an approximately constant flagella number per cell. This simple regulatory scheme ensures a fully motile population while keeping resource demands to synthesize and rotate flagella to a minimum. The findings reported here have implications for bacterial motility from the ecological perspective, particularly concerning its role in promoting fitness across different environments. Previous works have highlighted the upregulation as a fingerprint of a specific starvation response with motility triggered when nutrients run out (22, 24–26). In contrast, we here propose that at least a part of the upregulation of swimming in poorer growth conditions is not a starvation response *per se* but an obligatory regulation to maintain flagella numbers and swimming in diverse growth-supporting conditions as cell size changes. This picture is in line with observations that bacterial cells quickly stop swimming (9), actively brake motor rotation (41, 42), and even release their flagella upon entering starvation (43, 44). Notably, the maintenance of cellular motility in growth-supporting conditions enables cell population to rapidly expand into unoccupied nutrient rich territories, boosting overall population growth (9). The growth advantage of such a navigated range expansion relies on cells being motile across conditions, and a delayed onset of motility only in response to starvation would nullify the fitness advantage (9). Therefore, the efficient regulation of motility genes described here does not only minimizes the resources required to build and fuel the motility machinery, but it also supports fast navigated range expansion which further boost fitness (9, 21, 29).

The findings further provide a new perspective on the relation between cell-size and growth itself. Throughout the text, we have referred to the change in motility gene expression as an up-regulation in poor nutrient conditions. But this change can also be viewed as a down-regulation in nutrient replete conditions when cells grow fast. Given that the goal of the flagella regulatory system is to maintain the number of flagella per cell, we can view the decreased flagella expression at fast growth also as a consequence of increased cell size at fast growth. This view leads us to suggest a physiological rationale for *E. coli*’s choice of cell size at different growth rates. It is generally preferrable for bacterial cells to keep a small biomass (i.e., cell size) as it promotes efficient diffusive transport, fast nutrient uptake, and strong dispersal (45, 46). However, in favorable conditions allowing for rapid growth, the translational machinery per biomass is the most growth-limiting factor (47, 48) and making cell-size larger can be beneficial to alleviate this bottleneck: By increasing its size at fast growth, the cell effectively reduces the amount of flagella proteins that need to be synthesized, thus allowing more proteomic resources to be allocated towards ribosomes and other components of the translation machinery. Quantitatively, flagella proteins comprise ~3.0 % of the total protein mass in slow carbon-limited conditions and ~0.7 % in rich-defined medium (11). Thus, by increasing its cell size, *E. coli* manages to “save” 2.3 % of the proteome that would have otherwise been tied up in flagella synthesis. To put this amount in perspective, the entire set of biosynthesis enzymes saved when cells are provided with all amino acids and nucleotides is only ~11 % of the proteome (comparing the proteome composition of cells grown in rich-defined medium supplemented with glucose to those grown in glucose minimal medium). This saving accounts for a large share of the increase of growth rate from 1.0 1/*h* in glucose minimal medium to 1.8 1/*h* in rich-defined medium (11), based on the well-established linear relation between the ribosome content and growth rate, where every percent-of-proteome added to the protein synthesis machinery results in an ~0.06 1/*h* increase in growth rate (11, 23, 47). Thus a 2.3 % saving in proteome allocation to flagella synthesis would amount to a gain of ~0.14 1/*h* for growth in rich medium. In other words, had *E. coli* kept its size at that in poor nutrient condition, then it would suffer a 0.14 1/*h* reduction in growth rate (from the observed growth rate of 1.8 1/*h*) in rich medium just due to motility expression alone. This proteome resource savings by a change of cell size should be similarly applicable to other cellular processes which demand protein expression on a per-cell basis, including cell division and cell pole maintenance. Therefore, increasing cell size at fast growth is a simple and effective strategy to reduce competition for proteome resources at fast growth, for *E. coli* and likely many other fast growing bacterial species.

## Acknowledgement

We thank Matteo Mori, Chenhao Wu and Christina Ludwig for providing proteomic data, and Angela Dawson and Ekaterina Krasnopeeva for providing the pTOF24 plasmids carrying S219C and sticky *fliC*. Tomoya Honda acknowledges the JASSO long-term graduate fellowship award. Leonardo Mancini and Teuta Pilizota acknowledge the support of Cunningham Trust award ACC/KWF/PhD1. Work in the Hwa lab is supported by the NIH through grant R01GM109069 and by the NSF through grant MCB 1818384.

## Supplementary Figures

**Fig. S1:**
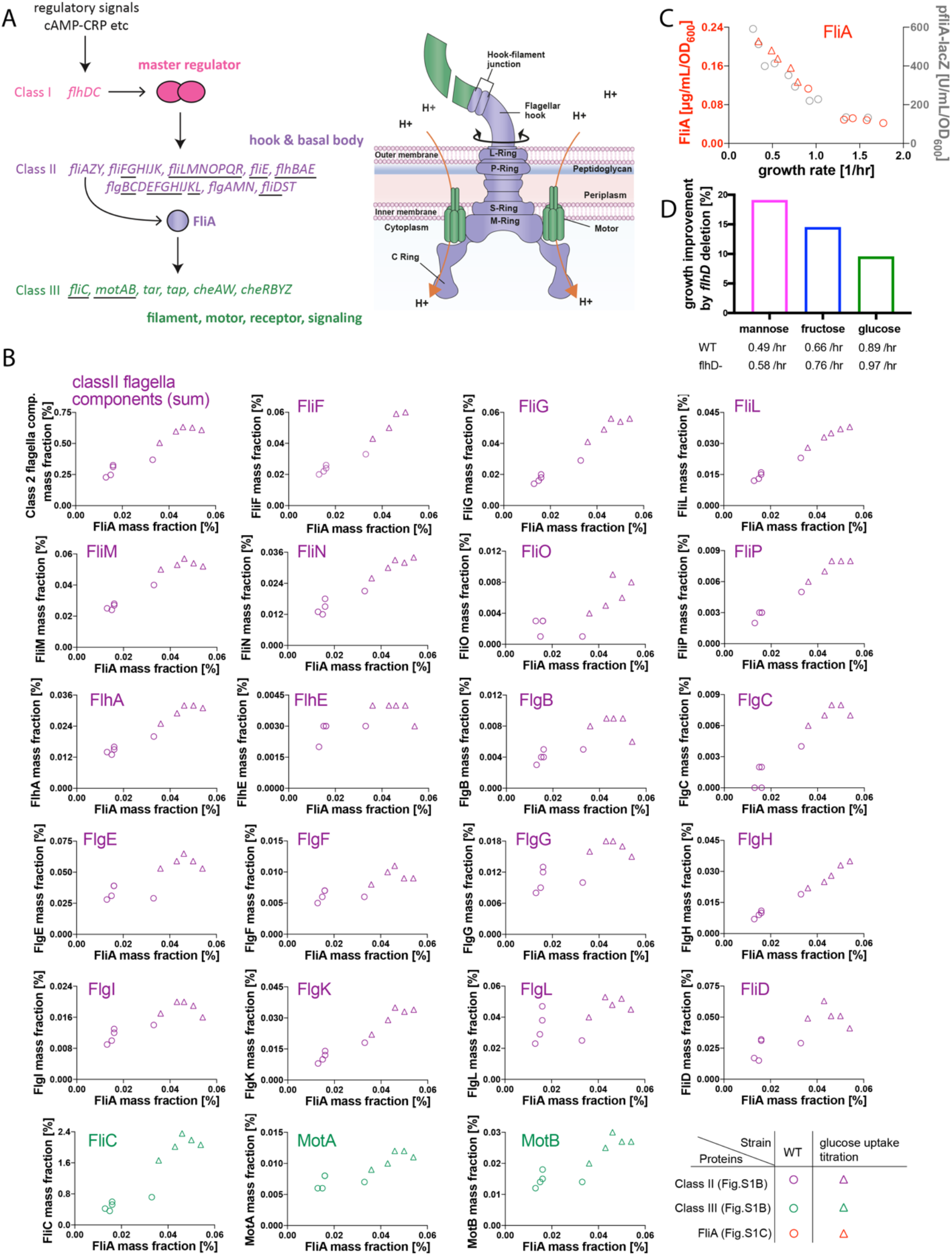
Overview of motility genes. (**A**) Schematic illustrations of motility gene regulation and flagella structure. The synthesis of flagella and chemotaxis systems requires more than 50 genes, encoded in multiple operons across the chromosome. Genes have been divided into three classes (30–32). In brief, the class I gene *flhDC* encodes a master regulator which drives the expression of class II genes that encode for hook-basal body elements (purple-colored) and a sigma factor *fliA*. FliA controls *fliC* and *motAB* expression which further encode for filament and motor structure components (green-colored), and other chemotactic machineries such as receptors and signal transduction. The rotation of the motor is powered by proton flux through the stator units (orange arrow). For the expression of the motility machinery, the master regulator, *flhDC*, requires a number of regulatory signals, including cAMP-CRP (51, 52). For simplicity, the diagram omits details of the regulation scheme, such as the anti-sigma factor activity of FlgM (53) and dual regulation of class II genes by both FlhDC and FliA (32). Physical components of flagella are underlined. **(B)** The mass fraction of class II (purple) and class III (green) proteins in relation to the mass fraction of the sigma-factor FliA (x-axis). The mass fraction is defined as mass of each protein divided by total protein mass in the sample. Most class II and class III proteins exhibit changes approximately proportional to FliA. Data were obtained from mass-spec measurements (11) quantifying the absolute protein abundance for the wild-type strain (NCM3722; circles) and for strains that allow the titration of glucose uptake (54) and thus growth rate (triangles; NQ1243 & NQ1390). For the wild-type, cells were grown in MOPS-based media with glucose (0.91 1/*h*), CAA (1.32 1/*h*), RDM (1.42 1/*h*), glucose+CAA (1.58 1/*h*), or glucose+RDM (1.77 1/*h*) as nutrient sources. The titratable strains NQ1243 and NQ1390 were grown with different inducer concentrations and glucose as a carbon source in M9-based media. **(C)** Comparison of protein abundance per biomass (from mass-spec data: red) and LacZ reporter expression (grey, same data as in Fig. 1A) for *fliA* which we used as a major reporter of class-II genes in this study. The agreement observed across growth rates together with the coordinated expression of class-II proteins with FliA in (B) supports the use of the P*fliA-lacZ* construct as a readout for the protein abundance of flagella hook-basal body elements. FliA abundance per biomass in unit of μ*g*/ *mL*/*OD*600 was obtained by multiplying FliA mass fraction by total protein mass per *mL* · *OD*600, which was characterized across the full range of growth conditions in Dai et al (55). (**D**) Growth rates of motility knockouts lacking the central regulator *flhD* (*flhD*-). The increases of growth rate relative to the WT (native *flhD*) are shown for different carbon sources (11 mM mannose, 20 mM fructose, 20 mM glucose). Growth rates for both, the WT (HE204) and the *flhD*- (HE275) strain are listed below. Growth measurements were repeated twice per condition and the averages are shown.

**Fig. S2:**
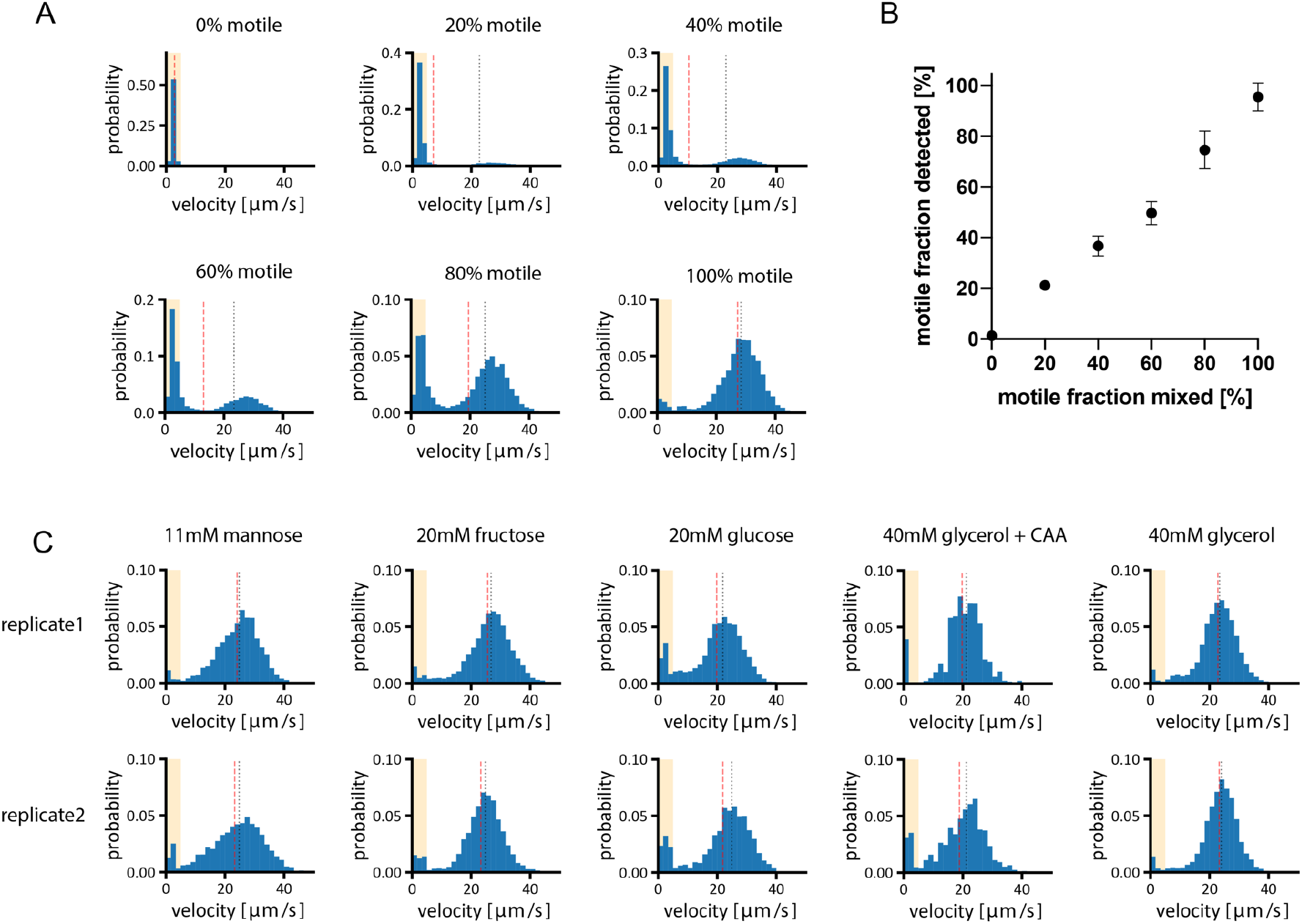
Cell to cell variation of swimming. (**A**) To establish our measurements of the motile fraction (**SI Text 1.3.3)**, we analyzed the cell to cell variation in swimming speeds observed for mixed populations consisting of different ratios of fully motile wild-type cells (HE206) and non-motile cells (HE275: Δ*flhD* strain). Shown are histograms obtained by the analysis of ~150 cells for each condition, with the mixing ratio shown on top of each histogram. When no motile cells are present (0 % motile), the mean velocity is 2.8 μm/s and, accordingly, we defined cells with a swimming velocity *v_i_* < 5 *μm/s* as non-motile (region highlighted in yellow). Red and grey lines indicate average swimming speed of all cells and motile cells only. In the shown experiment, both strains were grown in 40 mM glycerol as a carbon source. (**B**) Our computational tracking analysis agree well with the different mixing ratios, as confirmed by plotting the ratio computationally detected against the ratios we set experimentally. Error bars represent s.d. of technical replicates (N=2 at 0 % & 100 %, N=4 at 20 %, 40 %, 60 %, 80 %). (**C**) Reproducibility of our measurements with wild-type cells (HE206). To illustrate the reproducibility of motility measurements, the distributions of two biological replicates are shown for five different growth conditions (carbon sources as indicated).

**Fig. S3:**
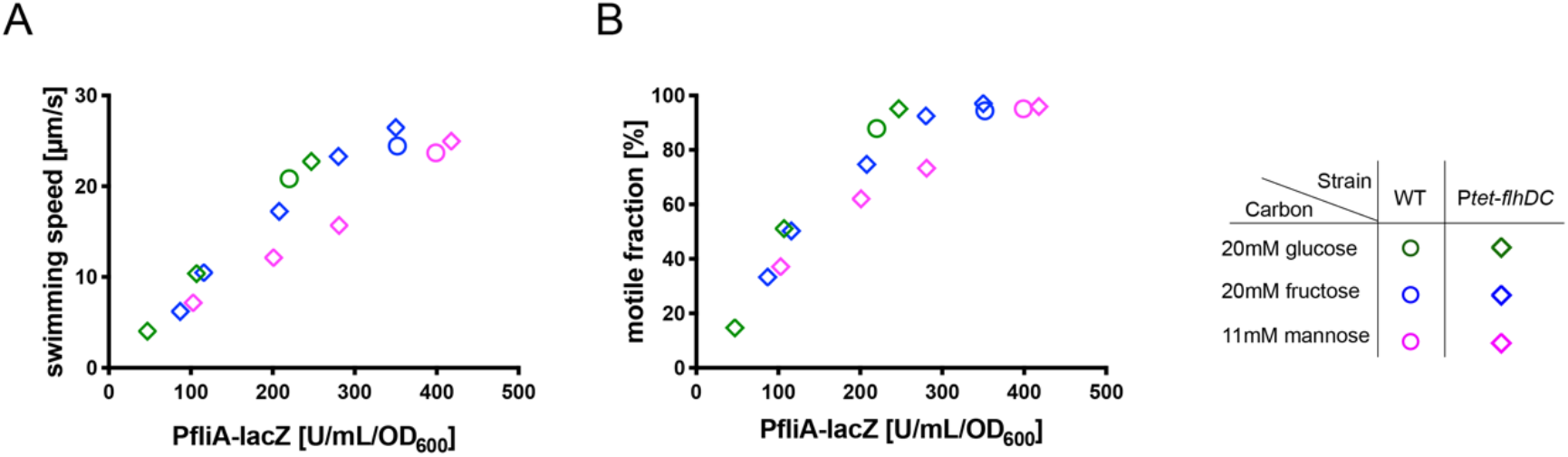
Relation between major swimming characteristics and the expression of *fliA* under *flhDC* titration. Average swimming speed **(A)** and the motile fraction (**B**) of cells against *fliA* expression, when grown in three different carbon sources with a titratable construct at different inducer levels (diamonds) as in Fig. 2A. The data of wild-type and titratable *flhDC* construct are shown as circles and diamonds, respectively. For experiments, strain HE206 and HE170 were used to obtain swimming characteristics, while HE207 and HE641 were used to obtain P*fliA-lacZ* expression data (the strains are identical except carrying different *lacZ* reporters; see **Table S1**).

**Fig. S4:**
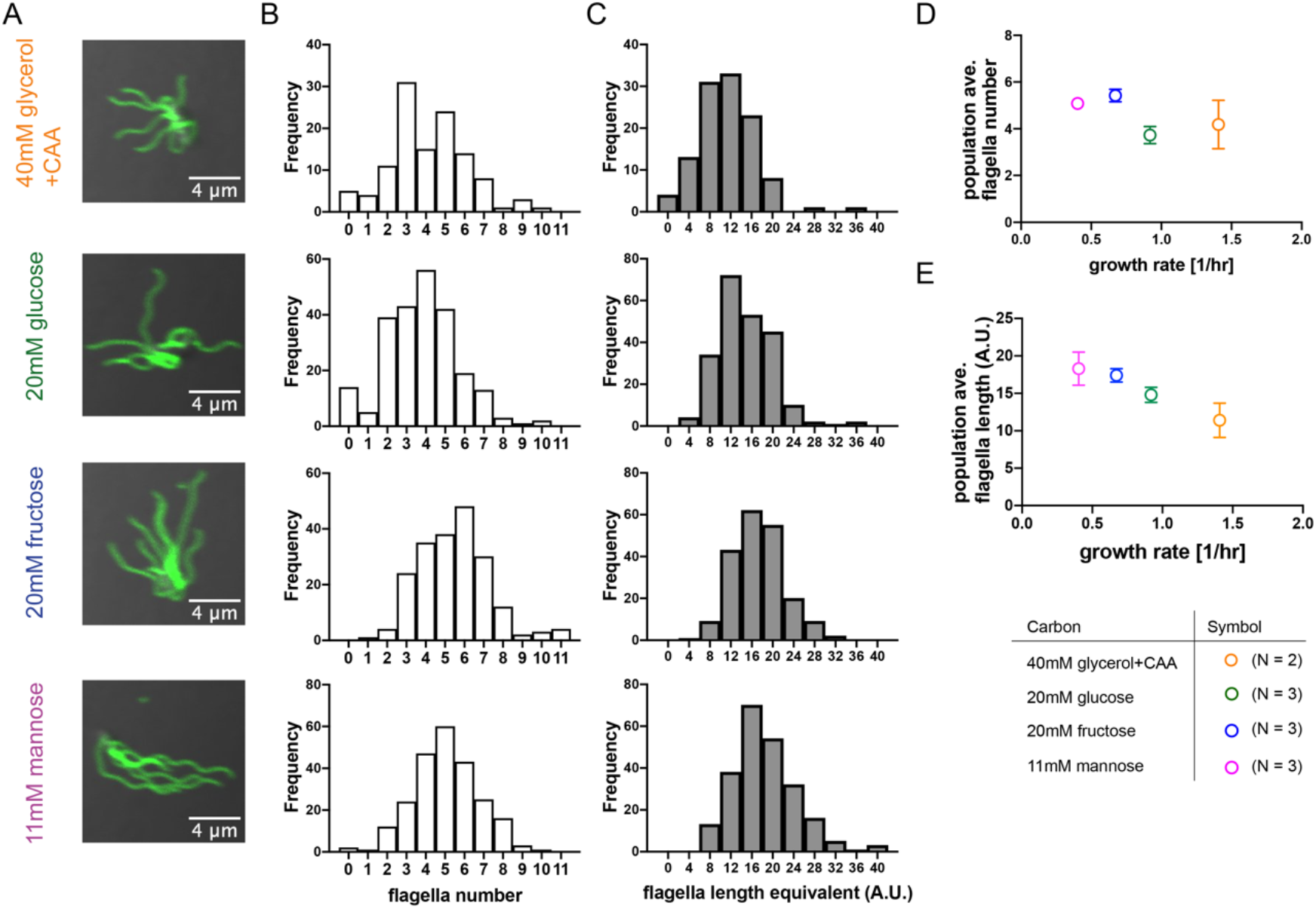
Variation of flagella filament number and length (native regulation of motility in WT background). To further analyze how the native regulation of motility genes relates to flagella number across conditions, we directly quantified the number and length of flagella filaments for cells growing on different carbon sources using a fluorescence staining assay. From top to bottom row, results for growth on glycerol+CAA (fastest condition), glucose, fructose, and mannose (slowest condition) are shown. **(A)** Flagella filament visualization. To visualize filament, we followed a previously established protocol (56) and used cells expressing a modified flagella filament component (S219C in *fliC*; strain HE582) which binds to the Alexa Fluor 488 Maleimide dye. Thus the filaments can be observed with a fluorescence microscope (57, 58) (**SI Text 1.5**). Representative images shown. **(B)** Variation of filament numbers per cell. By analyzing 60-100 of cells per each growth condition, we derived histograms for the filament number per cell. Notice that almost all cells have 2-7 filaments and that cells without flagella filaments are rarely present. **(C)** Variation of filament lengths. To obtain filament length we analyzed the fluorescence signal per filament (filament length equivalent) for different cells (details in **SI Text 1.5**). Both the number and length variation follow approximately a normal distribution. **(D)** and **(E)** Mean filament number per cell (D) and the length equivalent (E) for the different conditions when plotted against growth rate. The filament number is approximately constant for different growth rates, while filament is approximately 1.6-fold longer at slow (mannose) compared to fast growth (glycerol+CAA). Error bars in (D) and (E) represent s.d. based on N=2 or N=3 biological replicates as indicated in the legend.

**Fig. S5:**
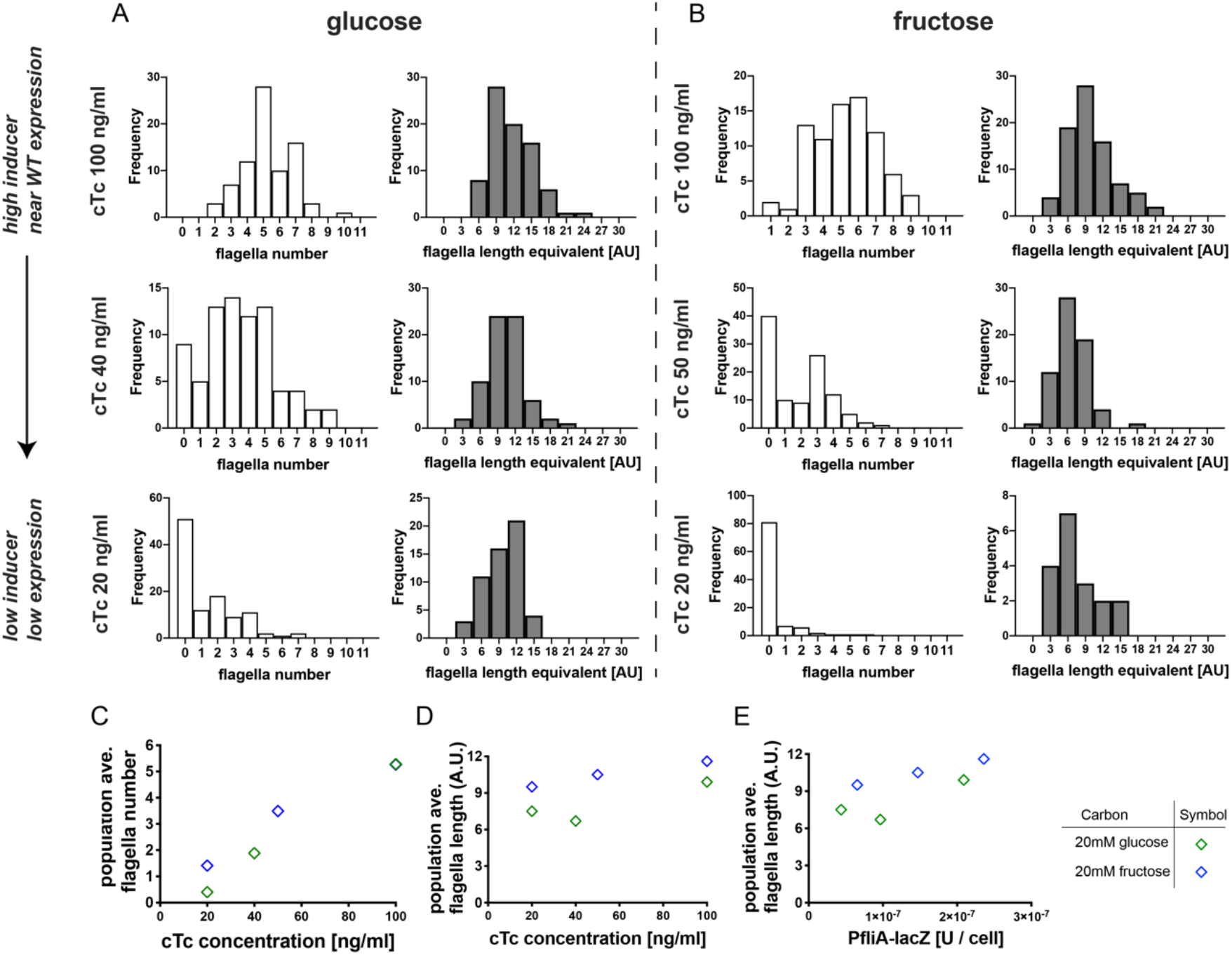
Variation of flagella filament number and length (*flhDC* titration). To further illustrate that the upregulation of motility genes at slow growth is required to maintain flagella filament number (and high motile fractions), we analyzed the variation of filament number and length when motility expression levels are changed by *flhDC* titration (strain construct as outlined in Fig. **2A**). **(A & B)** Histograms of filament number per cell and filament length equivalents are shown for growth in glucose (A) and fructose (B), which were determined via the fluorescence staining of flagella filament as described in **Fig. S**4 and **SI Text 1.5**. Strain HE571 was used for the experiments. We probed three different inducer levels (100, 40 or 50, and 20 ng/ml cTc as shown in top, middle, and bottom row). Notably, the fraction of cells with zero or only one filament is high when inducer levels are low (20 or 40 or 50 ng/ml cTc). In contrast, the variation of filament length is not changing much with inducer level. This can also be seen when mean filament number and length are plotted against inducer concentration in (**C**) and (**D**), or plotted against P*fliA-lacZ* expression per cell (**E**). Thus, the mean flagella filament number strongly changes with inducer concentrations, while mean length changes weakly.

## References

1. D. Molenaar, R. van Berlo, D. de Ridder, B. Teusink, Shifts in growth strategies reflect tradeoffs in cellular economics. Molecular Systems Biology 5, 323 (2009).

2. H. C. Berg, E. coli in Motion (Springer-Verlag, 2004) https://doi.org/10.1007/b97370 (April 19, 2021).

3. U. Alon, M. G. Surette, N. Barkai, S. Leibler, Robustness in bacterial chemotaxis. Nature 397, 168–171 (1999).

4. V. Sourjik, N. S. Wingreen, Responding to chemical gradients: bacterial chemotaxis. Curr Opin Cell Biol 24, 262–268 (2012).

5. A. J. Waite, N. W. Frankel, T. Emonet, Behavioral Variability and Phenotypic Diversity in Bacterial Chemotaxis. Annual Review of Biophysics 47, 595–616 (2018).

6. J. Adler, Chemotaxis in bacteria. Science 153, 708–716 (1966).

7. I. Chet, R. Mitchell, Ecological Aspects of Microbial Chemotactic Behavior. Annual Review of Microbiology 30, 221–239 (1976).

8. D. A. Koster, A. Mayo, A. Bren, U. Alon, Surface Growth of a Motile Bacterial Population Resembles Growth in a Chemostat. Journal of Molecular Biology 424, 180–191 (2012).

9. J. Cremer, et al., Chemotaxis as a navigation strategy to boost range expansion. Nature 575, 658–663 (2019).

10. D. W. Erickson, et al., A global resource allocation strategy governs growth transition kinetics of Escherichia coli. Nature 551, 119–123 (2017).

11. M. Mori, et al., From coarse to fine: The absolute Escherichia coli proteome under diverse growth conditions. in press Molecular Systems Biology.

12. D. C. Fung, H. C. Berg, Powering the flagellar motor of Escherichia coli with an external voltage source. Nature 375, 809–812 (1995).

13. C. V. Gabel, H. C. Berg, The speed of the flagellar rotary motor of Escherichia coli varies linearly with protonmotive force. PNAS 100, 8748–8751 (2003).

14. T. Minamino, Protein export through the bacterial flagellar type III export pathway. Biochimica et Biophysica Acta (BBA) - Molecular Cell Research 1843, 1642–1648 (2014).

15. E. J. Gauger, et al., Role of Motility and the flhDC Operon in Escherichia coli MG1655 Colonization of the Mouse Intestine. Infection and Immunity 75, 3315–3324 (2007).

16. X. Wang, T. K. Wood, IS 5 inserts upstream of the master motility operon flhDC in a quasi-Lamarckian way. The ISME Journal 5, 1517–1525 (2011).

17. M. Basan, et al., Overflow metabolism in Escherichia coli results from efficient proteome allocation. Nature 528, 99–104 (2015).

18. X. Yi, A. M. Dean, Phenotypic plasticity as an adaptation to a functional trade-off. eLife 5, e19307 (2016).

19. D. T. Fraebel, et al., Environment determines evolutionary trajectory in a constrained phenotypic space. Elife 6, e24669 (2017).

20. B. Ni, et al., Evolutionary Remodeling of Bacterial Motility Checkpoint Control. - PubMed - NCBI. Cell Reports 18, 866–877 (2017).

21. W. Liu, J. Cremer, D. Li, T. Hwa, C. Liu, An evolutionarily stable strategy to colonize spatially extended habitats. Nature 575, 664–668 (2019).

22. C. D. Amsler, M. Cho, P. Matsumura, Multiple factors underlying the maximum motility of Escherichia coli as cultures enter post-exponential growth. Journal Of Bacteriology 175, 6238–6244 (1993).

23. S. Hui, et al., Quantitative proteomic analysis reveals a simple strategy of global resource allocation in bacteria. Mol Syst Biol 11, e784–e784 (2015).

24. B. Ni, R. Colin, H. Link, R. G. Endres, V. Sourjik, Growth-rate dependent resource investment in bacterial motile behavior quantitatively follows potential benefit of chemotaxis. PNAS 117, 595–601 (2020).

25. M. Liu, et al., Global Transcriptional Programs Reveal a Carbon Source Foraging Strategy by Escherichia coli. J Biol Chem 280, 15921–15927 (2005).

26. K. Zhao, M. Liu, R. R. Burgess, Adaptation in bacterial flagellar and motility systems: from regulon members to ‘foraging’-like behavior in E. coli. Nucleic Acids Res 35, 4441–4452 (2007).

27. J. Saragosti, et al., Directional persistence of chemotactic bacteria in a traveling concentration wave. PNAS 108, 16235–16240 (2011).

28. X. Fu, et al., Spatial self-organization resolves conflicts between individuality and collective migration. Nature Communications 9, 2177 (2018).

29. S. Gude, et al., Bacterial coexistence driven by motility and spatial competition. Nature 578, 588–592 (2020).

30. G. S. Chilcott, K. T. Hughes, Coupling of Flagellar Gene Expression to Flagellar Assembly in Salmonella enterica Serovar Typhimurium and Escherichia coli. Microbiol Mol Biol Rev 64, 694–708 (2000).

31. S. Kalir, et al., Ordering Genes in a Flagella Pathway by Analysis of Expression Kinetics from Living Bacteria. Science 292, 2080–2083 (2001).

32. D. M. Fitzgerald, R. P. Bonocora, J. T. Wade, Comprehensive Mapping of the Escherichia coli Flagellar Regulatory Network. PLOS Genetics 10, e1004649 (2014).

33. W. S. Ryu, R. M. Berry, H. C. Berg, Torque-generating units of the flagellar motor of Escherichia coli have a high duty ratio. Nature 403, 444–447 (2000).

34. E. Krasnopeeva, C.-J. Lo, T. Pilizota, Single-Cell Bacterial Electrophysiology Reveals Mechanisms of Stress-Induced Damage. Biophysical Journal 116, 2390–2399 (2019).

35. J. H. Miller, Experiments in molecular genetics (1972) (May 11, 2021).

36. M. Basan, et al., Inflating bacterial cells by increased protein synthesis. Mol Syst Biol 11, 836–836 (2015).

37. C. L. Woldringh, J. S. Binnerts, A. Mans, Variation in Escherichia coli buoyant density measured in Percoll gradients. J Bacteriol 148, 58–63 (1981).

38. M. Schaechter, O. MaalØe, N. O. Kjeldgaard, Dependency on Medium and Temperature of Cell Size and Chemical Composition during Balanced Growth of Salmonella typhimurium. Microbiology, 19, 592–606 (1958).

39. F. Si, et al., Invariance of Initiation Mass and Predictability of Cell Size in Escherichia coli. Current Biology 27, 1278–1287 (2017).

40. H. Zheng, et al., General quantitative relations linking cell growth and the cell cycle in Escherichia coli. Nature Microbiology 5, 995–1001 (2020).

41. A. Boehm, et al., Second Messenger-Mediated Adjustment of Bacterial Swimming Velocity. Cell 141, 107–116 (2010).

42. K. Paul, V. Nieto, W. C. Carlquist, D. F. Blair, R. M. Harshey, The c-di-GMP Binding Protein YcgR Controls Flagellar Motor Direction and Speed to Affect Chemotaxis by a “Backstop Brake” Mechanism. Molecular Cell 38, 128–139 (2010).

43. J. L. Ferreira, et al., γ-proteobacteria eject their polar flagella under nutrient depletion, retaining flagellar motor relic structures. PLOS Biology 17, e3000165 (2019).

44. X.-Y. Zhuang, et al., Live-cell fluorescence imaging reveals dynamic production and loss of bacterial flagella. Molecular Microbiology 114, 279–291 (2020).

45. A. L. Koch, WHAT SIZE SHOULD A BACTERIUM BE? A Question of Scale. Annu. Rev. Microbiol. 50, 317–348 (1996).

46. K. D. Young, The Selective Value of Bacterial Shape. Microbiol. Mol. Biol. Rev. 70, 660–703 (2006).

47. M. Scott, C. W. Gunderson, E. M. Mateescu, Z. Zhang, T. Hwa, Interdependence of Cell Growth and Gene Expression: Origins and Consequences. Science 330, 1099–1102 (2010).

48. M. Scott, S. Klumpp, E. M. Mateescu, T. Hwa, Emergence of robust growth laws from optimal regulation of ribosome synthesis. Molecular Systems Biology 10, 747 (2014).

49. J. Rosko, V. A. Martinez, W. C. K. Poon, T. Pilizota, Osmotaxis in Escherichia coli through changes in motor speed. PNAS 114, E7969–E7976 (2017).

50. L. Mancini, et al., Escherichia coli’s physiology can turn membrane voltage dyes into actuators. bioRxiv, 607838 (2019).

51. K. A. Fahrner, H. C. Berg, Mutations That Stimulate flhDC Expression in Escherichia coli K-12. Journal of Bacteriology 197, 3087–3096 (2015).

52. O. Soutourina, et al., Multiple Control of Flagellum Biosynthesis in Escherichia coli: Role of H-NS Protein and the Cyclic AMP-Catabolite Activator Protein Complex in Transcription of the flhDC Master Operon. Journal of Bacteriology 181, 7500–7508 (1999).

53. K. Ohnishi, K. Kutsukake, H. Suzuki, T. Lino, A novel transcriptional regulation mechanism in the flagellar regulon of Salmonella typhimurium: an anti-sigma factor inhibits the activity of the flagellum-specific Sigma factor, σF. Molecular Microbiology 6, 3149–3157 (1992).

54. C. You, et al., Coordination of bacterial proteome with metabolism by cyclic AMP signalling. Nature 500, 301–306 (2013).

55. X. Dai, et al., Reduction of translating ribosomes enables Escherichia coli to maintain elongation rates during slow growth. Nature Microbiology 2, 1–9 (2016).

56. L. Turner, H. C. Berg, “Labeling Bacterial Flagella with Fluorescent Dyes” in Bacterial Chemosensing: Methods and Protocols, Methods in Molecular Biology., M. D. Manson, Ed. (Springer, 2018), pp. 71–76.

57. L. Turner, R. Zhang, N. C. Darnton, H. C. Berg, Visualization of Flagella during Bacterial Swarming. Journal of Bacteriology 192, 3259–3267 (2010).

58. L. Turner, A. S. Stern, H. C. Berg, Growth of Flagellar Filaments of Escherichia coli Is Independent of Filament Length. Journal of Bacteriology 194, 2437–2442 (2012).

