## Supplementary Text for "Coordination of gene expression with cell size enables *Escherichia coli* to efficiently maintain motility across conditions"

### Supplementary Information

TABLE OF CONTENTS

Supplementary Text 2
1. Experimental Method 2
1.1 Strain information 2
1.1.1 Major strain used in this study 2
1.1.2 Construction of P*fliA-lacZ* and P*fliC-lacZ* fusion 2
1.1.3 Construction of titratable *flhDC* strain 2
1.1.4 Strains for motor speed measurement and flagella staining 3
1.2 Media and growth conditions 3
1.2.1 Growth media for batch culture growth rate measurement 3
1.2.2 Strain culturing and growth measurement 3
1.3 Swimming behavior measurement 4
1.3.1 Swimming assay with capillary tube 4
1.3.2 Image analysis and statistical analysis 5
1.3.3 Calibration of motile cell detection 5
1.4 β-galactosidase assay 6
1.5 Flagella number and length quantification by staining 6
1.6 Motor speed measurement 7
1.6.1 Experimental procedure 7
1.6.2 Data analysis 7
2. Gene expression per biomass or cell-size 7
Supplementary Tables 9
Table S1: Strains used in this study. 9
Table S2: Growth rates and P*fliA-lacZ* expression. 10
Table S3: Swimming characteristics in wild-type strain. 10
Table S4: Motor rotation frequency in wild-type strain. 11
Table S5: P*fliA-lacZ* expression under *flhDC* titration. 11
Table S6: Swimming characteristics under *flhDC* titration. 12
Table S7: Flagella filament number and length in wild-type background 12
Table S8: Flagella filament number and length under *flhDC* titration 13
Table S9: primers used in this study. 13
Reference 14

### SUPPLEMENTARY TEXT

#### 1. Experimental Method

#### 1.1 Strain information

##### 1.1.1 Major strain used in this study

We use a motile variant of the *E. coli* K-12 (strain NCM3722B) whose physiology has been well-characterized in previous studies (1–5). Similar to other motile K-12 strains (6, 7), strain NCM3722B carries a 1kb insertion element (IS1) upstream of *flhDC* transcription site that activates *flhDC* expression and the motile phenotype (8, 9). The detailed information of the insertion element and the comparison to other motile K-12 strains are described in the Supplementary Text of our previous study (10).

##### 1.1.2 Construction of P*fliA-lacZ* and P*fliC-lacZ* fusion

Using primers PfliA-Xho-F/PfliA-BamH-R (**Table S9**), the *fliA* regulatory region (-313th nucleotide to the 1st nucleotide upstream of the *fliA* translational start point) was amplified from BW25113 genomic DNA. The PCR product was digested with *Xho*I and *BamH*I and then inserted into the corresponding sites of the plasmid pKDT (11) yielding the plasmids pKDT-P*fliA*. Present in this plasmid, the "*km*:*rrnB*T:P*fliA*" DNA fragment was amplified using primers PfliA-Z1/PfliA-Z2, and subsequentially integrated into the chromosome of *E.coli* strain EQ42 (11) to replace the region containing *lacI* and the entire *lacZ* promoter. The P*fliA-lacZ* fusion bearing kanamycin marker was confirmed by DNA sequencing. The constructed *PfliA-lacZ* was subsequently transferred into HE205 (NCM3722B) by phage P1*vir* transduction, resulting in strain HE207.
Similarly, we also prepared a lacZ reporter for P*fliC*. We substituted both *lacI* and *lacZ* promoter in EQ42 with the *fliC* regulatory region (-267th nucleotide to the 1st nucleotide upstream of the *fliC* translational start point). The constructed *PfliC-lacZ* was subsequently transferred into HE205 by phage P1*vir*, resulting HE206. The primers used for P*fliC* amplification and chromosomal integration are also listed in **Table S9.**

##### 1.1.3 Construction of titratable *flhDC* strain

Using the plasmid pKDT:Ptet (11) as template, the DNA fragment (referred to as "km:rrnBT:Ptet") containing the *km* gene, the *rrnB* terminator (rrnBT) and the Ptet promoter (including its 5'-UTR) was amplified using the primer pair Ptet.flh-P1/Ptet.flh-P2 (see **Table S9**). The PCR products were integrated into the chromosome of the K12-strain BW25113, which has no insertion element upstream of the *flhDC* regulatory region, using the method of Datsenko and Wanner (12) to replace the *flhDC* promoter: the upstream region from the 644$^{th}$ nucleotide to the 1$^{st}$ nucleotide relative to the translational start point of *flhD*. The chromosomal integration was confirmed first by colony PCR and subsequently by DNA sequencing. The region carrying "km:rrnBT:Ptet-*flhDC*" was transferred to the strains that carry P*tet-tetR* and P*fliC-lacZ* or P*fliA-lacZ* by phage P1*vir* transduction, yielding strains HE170 and HE641 respectively.

##### 1.1.4 Strains for motor speed measurement and flagella staining

For motor speed measurement, we replaced the native *fliC* sequence on the chromosome with a sticky *fliC* variant that renders the flagella filament sticky to glass surfaces and polystyrene beads (13, 14). The gene replacement was performed by following a protocol described in Merlin et al. (15), using a pTOF24 plasmid that carries the sticky *fliC* sequence (14). A sticky *fliC* strain (HE608) was created from our WT strain HE206, and the gene replacement was verified by sequencing. The experimental protocol to measure rotation frequencies is described in **Supplementary Text 1.6**.

A similar procedure was used to create a strain carrying the S219C *fliC* variant we used for flagella filament staining, using another pTOF24 plasmid that carries the S219C *fliC* sequence. The single substitution to cysteine enables direct labeling of flagella filament by sulfhydryl-specific Alexa Fluor maleimide dyes (16–18). The experimental protocol is described in **Supplementary Text 1.5**. HE582 (WT background) and HE571 (titratable *flhDC* background) were created from HE206 and HE301, respectively.

#### 1.2 Media and growth conditions

##### 1.2.1 Growth media for batch culture growth rate measurement

All growth media used in this study were based on the MOPS-buffered minimal medium used by Cayley et al. (19) with slight modifications. The base medium contains 40 mM MOPs and 4 mM tricine (adjusted to pH 7.4 with NaOH), 1.32 mM $KH_2PO_4$, 0.523 mM $MgCl_2$, 0.276 mM $Na_2SO_4$, 0.1 mM $FeSO_4$, 0.1 M NaCl and 20 mM $NH_4Cl$ as nitrogen source. The trace micronutrients were not added into the MOPs medium, since the metal components were reported to inhibit motility of *E. coli* (20). For growth measurements in minimum medium, one of the followings was used as the primary carbon source: 20 mM glucose, 10 mM maltose, 20 mM fructose, 40 mM glycerol, 20 mM galactose, 20 mM sorbitol, 11 mM mannose, 5 mM mannose, 20 mM aspartate. For rich growth conditions, the following media were used: 40 mM glycerol minimum medium with 2 mM of nine amino acids (aspartate, serine, histidine, isoleucine, leucine, lysine, methionine, phenylalanine, valine), 40 mM glycerol minimum medium with 2% casamino acid (CAA: BD 223050), and 20 mM glucose minimum medium with RDM (Rich Defined Medium: Teknova M2103 & M2104). For the strains titratable for *flhDC* (HE170, HE571), different concentrations of chlortetracycline (cTc) were additionally provided into the medium.

For the swimming assay and the flagella staining (details below), 0.05 % PVP40 (Sigma-Aldrich: polyvinylpyrrolidone average mol 40,000) was additionally provided into the medium. PVP40 prevents cells and flagella from binding to material surfaces and greatly reduces handling variation when running the assay (21).

##### 1.2.2 Strain culturing and growth measurement

Cells were grown in a 37°C water bath shaker shaking at 250 rpm. The culture volume was no more than 4.5 ml in 18 mm × 150 mm test tubes (Fisher Scientific) to limit the depth of the culture in tubes for aeration purpose. For growth in minimum medium, growth experiment was carried out in three steps: “seed culture” in LB broth, “pre-culture” and “experimental culture” in identical minimal medium. For seed culture, one colony from fresh LB agar plate was inoculated into liquid LB and cultured at 37°C with shaking. After 4-5 hours, cells were centrifuged and washed once with the desired minimal medium. Cells were then diluted into the minimal medium and cultured in the 37°C water bath shaker overnight (pre-culture). The starting $OD_{600}$ in the pre-culture was adjusted so that exponential cell growth was maintained throughout overnight incubation, preventing cells from reaching saturation. Cells from the overnight pre-culture were then diluted to $OD_{600}$ = 0.005-0.02 in pre-warmed minimal media and cultured again in the 37°C water bath shaker (experimental culture). After cells were grown at least for three generations, $OD_{600}$ was measured around every half doubling of cell growth. At each time point, $OD_{600}$ was measured by collecting a sample volume of 200 µl cell culture in a cuvette (Starna Cells, Atascadero, CA) and using a spectrophotometer (Thermo Scientific). 4-6 $OD_{600}$ data points within the range 0.04 to 0.3 were used for calculating growth rate.

For growth in rich conditions, it was difficult to maintain exponential cell growth during overnight pre-culture. Therefore, the saturated overnight culture was diluted into fresh medium and continuously grown for about 10 generations. During this period, the bacterial density was always maintained below $OD_{600}$ = 0.4, by repeating growth and dilution. The culture was then diluted into fresh medium with an $OD_{600}$ below 0.01 to start the experimental culture. Note that in our previous study (10), we found maintaining exponential growth is important to avoid low motility due to outgrowth from stationary phase and to capture swimming behavior during balanced exponential growth.

#### 1.3 Swimming behavior measurement

##### 1.3.1 Swimming assay with capillary tube

During steady state growth ($OD_{600}$ = 0.05-0.3), samples were taken and analyzed for swimming behavior. For each time point, a sample volume of about 200 µl was taken from the batch culture and filtered. The filtered medium (free of bacteria) and a calculated volume from the batch culture were then used to prepare a cell-culture sample with a cell density diluted down to $OD_{600} \approx 0.005$. Sampling and dilution were done using pre-heated Eppendorf tubes and filters (drybath, 37°C). This dilution was chosen to prevent difficulty in cell-tracking at high cell densities while keeping cell numbers high enough to ensure good statistics. The dilution with filtered media prevents cells from encountering strong environmental shifts before the quantification of swimming behavior. Immediately after dilution, one end of a pre-heated 300 µm glass capillary tube (rectangular shape, VitroCom) was then inserted into the diluted culture. Following capillary forces, the culture was filled the capillary within a few seconds. The width of the capillary minimized interactions of cells with the glass surfaces. The completely filled capillary was sealed with silicone grease (to prevent fluid flow) and immediately placed under a microscope (Nikon TI-U) covered with a chamber (InVivo Scientific) maintained at 37 °C. Cells were observed using a 10x phase-contrast objective with the focus centered between the upper and lower glass-surface of the capillary. Images were acquired using a CCD camera (EO-13122M Edmund Optics). Movies of swimming cells were recorded at 20 frames per second for a total length of up to 150 s. 100 to 300 cells were detected in each movie. For each growth condition, two to three movies were recorded at different cell densities in the culture ($OD_{600}$) during steady state growth. The reported values of swimming speed and motile fraction are the averages of measurements at different $OD_{600}$.

##### 1.3.2 Image analysis and statistical analysis

To obtain swimming characteristics, the obtained movies were analyzed in three consecutive steps using a custom-made Python script: 1. cell-detection, 2. cell-tracking over time, 3. trajectory analysis. The script is available online at: https://github.com/jonascremer/swimming_analysis.

1) Cell-detection: Cells within each frame were detected using an adaptive thresholding algorithm. First, background intensity was removed in two ways, either by subtracting the intensity averaged over a time-window of 200 frame (10 seconds), or by performing an adaptive background subtraction for which the intensities of neighboring pixels (100px) were averaged and subtracted for each time-step. The latter allowed to also detect non-moving cells. Cells were then detected by thresholding with the threshold set at 50% of the maximum intensity observed. Connected areas remaining after thresholding were each counted as one cell and the location of each cell was determined by weighting the different pixel positions with intensity.

2) Cell-tracking: To track cells over time, determined cell positions were linked from frame to frame following the algorithm developed by Crocker and Grier (22) utilizing the Python package trackpy (https://soft-matter.github.io/trackpy). For each sample analyzed, detection and trajectory generation were checked manually (by observing generated movies were detected cell positions and generated trajectories were shown on top of the original images obtained).

3) Trajectory analysis: Swimming behavior was quantified by a statistical analysis of the obtained trajectories. To this end, each trajectory (j) was first analyzed for tumbling events ($i^j$) defined by large changes in angle or velocity: Instantaneous changes in velocity and angle were calculated for each time-point using averages over 4 frames. Instantaneous speed changes of more than 50 % between frames or a change in angle $|\Delta\theta| > 1.2$ were assigned to tumble events. The velocity $v_j$ for each trajectory was obtained by calculating the spatial displacement (in two dimensions: $dR^j = \sum_i dr_i^j = \sqrt{{dx_i^j}^2 + {dy_i^j}^2}$) and the total time ($dT^j = \sum_i dt_i^j$) when cells were not tumbling but running $\{dt_i\}$: $v_j = dR^j / dT^j$. Running time was obtained by the average time between observed tumbling events. To obtain the averages and standard deviation of swimming speed and running time observed across the whole sample, the obtained values for each trajectory were weighted with the length of the trajectory.

##### 1.3.3 Calibration of motile cell detection

To distinguish motile from non-motile cells in our analysis, we conducted a calibration experiment by mixing motile and non-motile cells (**Fig. S2**). Specifically, we grew wild-type cells (HE206) and non-motile cells (HE275: Δ*flhD* strain), respectively, in glycerol as a carbon source. When cell cultures reached around $OD_{600} \approx 0.2$, they were diluted down to $OD_{600} \approx 0.005$ as described in **Supplementary Text 1.3.1**. Then, the diluted cultures were mixed at different ratios and swimming characteristics of the population was quantified. We prepared multiple technical replicates for each mixing ratio. When only non-motile cells were present (Δ*flhD* strain), cells moved with an average dislocation speed of only 2.8 μm/s (0.78 μm/s standard deviation) as shown in **Fig. S2**. In contrast, swimming cells move with much higher velocities. Hence, we set 5 μm/s as a threshold to distinguish between motile and non-motile cells. This approach successfully recovers the fraction of motile cells provided for the different mixing ratios (**Fig. S2**).

#### 1.4 β-galactosidase assay

The assay was performed following a similar protocol as detailed in a previous study (2). In brief, samples (0.2 ml cell culture) were collected, fast frozen on dry ice and stored at -80 °C prior to running the β-galactosidase assay. About four samples were collected for each culture during exponential growth (for $OD_{600}$ = 0.1~0.4). For each sample collected, β-galactosidase activity was measured at 37 °C by a traditional Miller method. The LacZ activity obtained (in units of U/ml or $OD_{420}$/min/ml) were plotted against the respective $OD_{600}$, and the resulting slope from a linear regression was taken to be the "LacZ expression level" (in units of U/ml $OD_{600}$).

#### 1.5 Flagella number and length quantification by staining

We used two strains, HE582 (WT background) and HE571 (titratable *flhDC* background), carrying the modification S219C in the *fliC* sequence (construction details provided in **Supplementary Text 1.1.4**). This amino acid substitution enables the direct labeling of flagella filaments by sulfhydryl-specific Alexa Fluor maleimide dyes (16–18). For staining, we collected cells during exponential growth when the cell cultures reached about $OD_{600} \approx 0.2$. 5ml culture was collected and the medium was removed after centrifuging at 1200 x *g* for 6 minutes at 37 °C. The cell pellet was gently resuspended by 500 µl pre-warmed motility buffer (0.01 M potassium phosphate at pH 7.0, $10^{-4}$ M EDTA, 0.067 M NaCl and 0.0001 % Tween 20). Flagella filaments were then labeled by adding the Alexa Fluor 488 Maleimide dye (ThermoFisher: A10254) at a final concentration 50 µg/ml and incubating the culture under dark conditions at 37 °C for 15 minutes. To get rid of the remaining free dye, 10 ml of pre-warmed motility buffer was then added, and the cell culture was centrifuged again at 1200 x *g* for 6 minutes at 37 °C. Cells were then gently resuspended in 5ml motility buffer. We further realized that growing cells in medium with 6.25 µg/ml dye during the main culturing step also provides sufficient levels of staining without impacting growth. This approach yielded similar results as the above centrifugation method with a much faster processing time and was used as well.

For imaging, 2.5 µl of cell culture was placed onto a cover glass and covered by a 1 cm x 1cm of 2 % agar pad. Cells were observed using a laser-scanning microscope (Leica SP8 inverted microscope equipped with 8Hz resonant scanner and HyD detector). Fluorophores were excited with a 488 nm laser line and the detectors scanned in the wave-length range 500−550 nm. Detection and laser intensity settings were kept constant throughout experiments preventing saturation of the intensity signal. During observations, samples were maintained at 37 °C within an environmental chamber.

The images were taken with an objective covering a size of 58.18 x 58.18 $\mu m^2$ containing 1024 x 1024 pixels. To reduce noise, each image was taken by using a 64x line-average scanning mode. Simultaneously, images were also taken with a transmitted light detector allowing the detection of cells without fluorescent signals. About 10 images were taken at different positions to analyze 60-100 cells in each replicate. We confirmed that the cells with wild-type *fliC* sequence exhibit little signal that is indistinguishable to background, thus the main fluorescent signals come from the stained filaments, not from the cell body.

The number of flagella filaments was counted manually. To ensure that all cells are considered, including those without any (stained) flagella, cells and their position were detected by looking at the corresponding transmission images which do not depend on fluorescent signals. To analyze filament length, images were processed by ImageJ (Fiji). First, an area around each cell covering all filaments was selected by the polygon tool. Then, integrated fluorescent intensity was measured for the selected areas. To obtain length equivalent values, the integrated intensity from each cell was divided by the number of filaments that the cell harbored. Since we noted some difference in staining levels depending on the media used, we further analyzed the mean fluorescence intensity observed along narrow sections of defined size along some filaments (3.01 x 0.57 μm) in each condition. Using these values, we corrected the intensity values for entire cells in each condition.

#### 1.6 Motor speed measurement

##### 1.6.1 Experimental procedure

Cells (sticky-*fliC*, strain HE608) were harvested from steady state cultures and flagella were sheared by passing the cells through two syringes with narrow-gauge needles (26 gauge) connected by plastic tubing. Sheared cells were then spun down at 8000 G for 2 minutes and the pellet resuspended in half the initial volume. Flow cells, as described in Mancini et al. (23), were flushed with PolyL-Lysine and immediately washed with roughly 10 flow chamber volumes (2.5 ml) of water and 1 ml of growth medium. Cells were then loaded in the slide and incubated for 10 minutes. Cells that did not stick to the surfaces of the chamber were washed out with 1 ml of growth medium and a suspension of beads with diameter 0.5 μm (Polysciences, US) was flown in the chamber and incubated for 10 minutes. Beads in excess were washed away with 1 ml of growth medium and the slides were moved to a pre-warmed microscope stage. The preparation and measurement procedure were carried out at 37 C and all the liquids had been pre-warmed. The motor speed was estimated via back-focal plane interferometry as explained previously (24). During the measurement, a continuous flow of growth medium was maintained using a peristaltic pump (Fusion 400, Chemyx, USA) at a flowrate of 100 μl per minute. Care was taken to ensure measurements in each slide were performed in a time window so that there is no movement of motor positions due to cell growth or no significant filament regrowth.

##### 1.6.2 Data analysis

Motor speed traces were analyzed by a moving-window discrete Fourier transform that moves in 100 ms intervals and has a size of 1.024 s. Values 45 and 55 and between -55 and -45 were discarded. The remaining values were taken as absolute and the median was calculated. Each data point is the median of the data points for one single cell. The reported values (average and standard deviation) in **Fig. 1E** are based on the analysis of 30-50 cells within each growth condition.

### 2. Gene expression per biomass or cell-size

To interpretate gene expression data, it is important to consider how expression levels are measured. Many common approaches quantify gene expression levels "*per biomass*". For LacZ and GFP reporter measurements in batch culture experiments, for example, the expression levels are typically normalized by volume and optical density ($OD_{600} \cdot ml$), a good proxy for biomass. Similarly, most omics approaches report the relative expression levels such as the fraction of a specific mRNA compared to total mRNA mass (transcriptomics) or the fraction of a specific protein compared to the entire protein mass (proteomics). By considering the total mRNA or protein content per biomass, the expression levels of genes of interest per biomass are obtained. In contrast, typical microscopy methods using fluorescence reporters give a readout of expression "*per cell*". When the (average) cell-size does not change much, the single-cell observations resemble the batch measurements; the fold change of gene expression for different conditions is similar whether gene-expression is quantified either *per cell* or *per biomass*. However, when cell-size changes with conditions, as being observed for *E. coli* when growth conditions change, one has to account for these changes to compare per cell and per biomass expression measurements. For example, to obtain the expression per cell and relate it to the number of flagella per cell, we here converted the gene-expression *per biomass* quantities from the batch culture measurements to *per cell* quantities using available cell-size measurements.

### SUPPLEMENTARY TABLES

**Table S1: Strains used in this study.**

| Strain | Parent | Genotype | Description | Ref. |
|---|---|---|---|---|
| NQ122 | NCM3722 | P*lacZ*::*Km*-P*tet-lacZ* | Strain carrying *lacZ* driven by constitutive P*tet* promoter | (2) |
| NQ840 | NCM3722 | *ycaD*::*rrnB*T-P*tet-tetR* | Strain carrying P*tet* promoter driven *tetR* at the *ycaD* site | (2) |
| HE136 | NQ840 | *ycaD*::*rrnB*T-P*tet-tetR* *Km*::P*fliC-lacZ* | NQ840 carrying P*fliC-lacZ* fusion. | This study |
| HE161 | HE136 | *ycaD*::*rrnB*T-P*tet-tetR* FRT::P*fliC-lacZ* | HE136 with *Km* gene flipped out. | This study |
| HE170 | HE161 | *ycaD*::*rrnB*T-P*tet-tetR* FRT::P*fliC-lacZ* *Km*:T:P*tet-flhDC* | HE161 carrying *flhDC* driven by P*tet* promoter and wild-type *fliC*. | This study |
| HE204 | NCM3722 | Δ*tcyJ*::*km* | *tcyJ* null NCM3722 carrying IS1 upstream of *flhDC* promoter and wild-type *fliC*. | (10) |
| HE205 | HE204 | Δ*tcyJ*::FRT | HE204 with *km* gene flipped out. | (10) |
| HE206 | HE205 | Δ*tcyJ*::FRT *Km*::P*fliC-lacZ* | HE205 carrying P*fliC-lacZ* fusion. | (10) |
| HE207 | HE205 | Δ*tcyJ*::FRT *Km*::P*fliA-lacZ* | HE205 carrying P*fliA-lacZ* fusion. | This study |
| HE275 | HE205 | Δ*tcyJ*::FRT Δ*flhD*::*km* | *flhD* deletion in HE205. Δ*flhD* is derived from CGSC11772 | This study |
| HE301 | HE170 | *ycaD*::*rrnB*T-P*tet-tetR* FRT::P*fliC-lacZ* FRT:P*tet-flhDC* | HE170 with *Km* gene flipped out. | This study |
| HE571 | HE301 | *ycaD*::*rrnB*T-P*tet-tetR* FRT::P*fliC-lacZ* FRT:P*tet-flhDC* *fliC* with S219C | HE301 carrying S219C in *fliC* gene. Used for staining. | This study |
| HE582 | HE206 | Δ*tcyJ*::FRT *Km*::P*fliC-lacZ* *fliC* with S219C | HE206 carrying S219C in *fliC* gene. Used for staining. | This study |
| HE608 | HE206 | Δ*tcyJ*::FRT *Km*::P*fliC-lacZ* Sticky *fliC* | HE206 carrying sticky *fliC* gene. Used for motor speed measurement. | This study |
| HE641 | | *ycaD*::*rrnB*T-P*tet-tetR* FRT::P*fliA-lacZ* *Km*:T:P*tet-flhDC* | Similarly constructed as HE170 except carrying *fliA-lacZ* reporter. | This study |

**Table S2: Growth rates and P*fliA-lacZ* expression.**

| Strain | Carbon source | Supplement | Growth rate (1/hr)[*] | P*fliA-lacZ* (U/ml/$OD_{600}$)[*] | P*fliA-lacZ* (U/cell)[+] |
|---|---|---|---|---|---|
| HE207 | 20 mM aspartate | | 0.28 | 591 | $2.46 \times 10^{-7}$ |
| | 20 mM fructose | | 0.68 | 352 | $2.37 \times 10^{-7}$ |
| | 20 mM glucose | | 0.93 | 220 | $1.97 \times 10^{-7}$ |
| | 5 mM mannose | | 0.34 | 512 | $2.30 \times 10^{-7}$ |
| | 10 mM maltose | | 0.76 | 293 | $2.15 \times 10^{-7}$ |
| | 20 mM sorbitol | | 0.53 | 413 | $2.30 \times 10^{-7}$ |
| | 11 mM mannose | | 0.41 | 399 | $1.95 \times 10^{-7}$ |
| | 40 mM glycerol | 2 mM of nine amino acids | 1.02 | 228 | $2.29 \times 10^{-7}$ |
| | 40 mM glycerol | 2 % casamino acids | 1.34 | 135 | $1.98 \times 10^{-7}$ |
| | 20 mM glucose | rich defined medium (RDM) | 1.60 | 133 | $2.63 \times 10^{-7}$ |

[*]All measurements were conducted once.
[+]To calculate the expression per cell, the expression per biomass (**U/ml/$OD_{600}$**) were multiplied by cellular mass (**$OD_{600}$·ml/cell**) expected from the fitting analysis in (Fig. 4B).

**Table S3: Swimming characteristics in wild-type strain.**

| Strain | Carbon source | Supplement | Growth rate (1/hr) | Motile fraction (%) | Swim speed (μm/s)[+] | n[*] |
|---|---|---|---|---|---|---|
| HE206 | 20 mM aspartate | | 0.25 | 90.5 | $20.7 \pm 9.9$ | 1 |
| | 20 mM fructose | | 0.67 | 94.4 | $24.4 \pm 8.0$ | 2 |
| | 20 mM glucose | | 0.92 | 87.9 | $20.8 \pm 9.2$ | 2 |
| | 11 mM mannose | | 0.40 | 95.1 | $23.7 \pm 8.7$ | 2 |
| | 40 mM glycerol | | 0.68 | 97.4 | $24.2 \pm 6.8$ | 3 |
| | 5 mM mannose | | 0.30 | 94.2 | $24.3 \pm 10.3$ | 1 |
| | 40 mM glycerol | 2 mM of nine amino acids | 1.16 | 89.8 | $19.8 \pm 8.2$ | 1 |
| | 40 mM glycerol | 2 % casamino acids | 1.41 | 90.9 | $19.3 \pm 7.9$ | 2 |
| | 20 mM glucose | rich defined medium (RDM) | 1.56 | 84.1 | $16.2 \pm 9.1$ | 1 |

[+]The values indicate mean and standard deviation of swimming speed in the population.
*The number of biological repeats used for swim measurements is shown.

**Table S4: Motor rotation frequency in wild-type strain.**

| Strain | Carbon source | Growth rate (1/hr) | Motor rotation frequency (Hz)* |
|---|---|---|---|
| HE608 | 20 mM fructose | 0.73 | $211 \pm 35$ |
| | 20 mM glucose | 0.87 | $231 \pm 32$ |
| | 11 mM mannose | 0.39 | $193 \pm 42$ |

*The values indicate mean and standard deviation of rotation frequency in the population. All measurements were conducted once.

**Table S5: P*fliA-lacZ* expression under *flhDC* titration.**

| Strain | Carbon source | cTc concentration (ng/ml) | Growth rate (1/hr)* | P*fliA-lacZ* (U/ml/$OD_{600}$)* | P*fliA-lacZ* (U/cell)*+ |
|---|---|---|---|---|---|
| HE641 | 20 mM glucose | 20 | 0.96 | 47 | $4.4 \times 10^{-8}$ |
| | | 40 | 0.94 | 107 | $9.7 \times 10^{-8}$ |
| | | 100 | 0.88 | 247 | $2.1 \times 10^{-7}$ |
| | 20 mM fructose | 20 | 0.77 | 87 | $6.5 \times 10^{-8}$ |
| | | 30 | 0.76 | 116 | $8.6 \times 10^{-8}$ |
| | | 50 | 0.73 | 208 | $1.5 \times 10^{-7}$ |
| | | 70 | 0.71 | 280 | $1.9 \times 10^{-7}$ |
| | | 100 | 0.69 | 350 | $2.4 \times 10^{-7}$ |
| | 11 mM mannose | 20 | 0.51 | 103 | $5.7 \times 10^{-8}$ |
| | | 30 | 0.49 | 201 | $1.1 \times 10^{-7}$ |
| | | 50 | 0.45 | 281 | $1.4 \times 10^{-7}$ |
| | | 100 | 0.40 | 418 | $2.0 \times 10^{-7}$ |

*All measurements were conducted once.
+To calculate the expression per cell, the expression per biomass (**U/ml/$OD_{600}$**) were multiplied by cellular mass (**$OD_{600}$·ml/cell**) following the exponential fit in Fig. 4B.

**Table S6: Swimming characteristics under *flhDC* titration.**

| Strain | Carbon source | cTc concentration (ng/ml) | Growth rate (1/hr) | Motile fraction (%) | Swim speed all cells (μm/s)+ | Swim speed motile cells (μm/s)* |
|---|---|---|---|---|---|---|
| HE170 | 20 mM glucose | 20 | 0.95 | 15 | 4.0 ± 4.5 | 13.4 ± 5.9 |
| | | 40 | 0.95 | 51 | 10.4 ± 8.9 | 17.5 ± 7.1 |
| | | 100 | 0.87 | 95 | 22.7 ± 7.9 | 23.8 ± 6.5 |
| | 20 mM fructose | 20 | 0.76 | 33 | 6.2 ± 5.7 | 13.5 ± 5.1 |
| | | 30 | 0.74 | 50 | 10.5 ± 8.9 | 17.8 ± 6.7 |
| | | 50 | 0.69 | 75 | 17.2 ± 9.2 | 21.8 ± 6.4 |
| | | 70 | 0.71 | 92 | 23.3 ± 10.0 | 25.0 ± 8.5 |
| | | 100 | 0.66 | 97 | 26.5 ± 7.5 | 27.2 ± 6.2 |
| | 11 mM mannose | 20 | 0.47 | 37 | 7.2 ± 6.4 | 14.7 ± 5.9 |
| | | 30 | 0.46 | 62 | 12.1 ± 8.7 | 17.6 ± 6.7 |
| | | 50 | 0.43 | 73 | 15.7 ± 9.7 | 20.4 ± 6.7 |
| | | 100 | 0.38 | 96 | 25.0 ± 9.8 | 25.9 ± 8.8 |

+The values indicate mean and standard deviation of swimming speed in the population.
*The values indicate mean and standard deviation of swimming speed in the motile fraction defined as $\boldsymbol{v_i}$ > $\mathbf{5}\ \mu\boldsymbol{m/s}$.
All measurements were conducted once.

**Table S7: Flagella filament number and length in wild-type background.**

| Strain | Carbon source | Supplement | Repeat number | Filament number* | Filament length equivalent (AU)* |
|---|---|---|---|---|---|
| HE582 | 20 mM glucose | | 3 | 3.7 ± 0.4 | 14.8 ± 1.0 |
| | 11 mM mannose | | 3 | 5.1 ± 0.2 | 18.3 ± 2.2 |
| | 20 mM fructose | | 3 | 5.4 ± 0.3 | 17.4 ± 0.9 |
| | 40 mM glycerol | 2 % casamino acids | 2 | 4.2 ± 1.0 | 11.4 ± 2.3 |

*The values indicate mean and standard deviation of biological replicates.

**Table S8: Flagella filament number and length under *flhDC* titration.**

| Strain | Carbon source | cTc concentration (ng/ml) | Filament number* | Filament length equivalent (AU)* |
|---|---|---|---|---|
| HE571 | 20 mM glucose | 20 | 0.4 | 7.5 |
| | | 40 | 1.9 | 6.7 |
| | | 100 | 5.3 | 9.9 |
| | 20 mM fructose | 20 | 1.4 | 9.5 |
| | | 50 | 3.5 | 10.5 |
| | | 100 | 5.3 | 11.6 |

*All measurements were conducted once.

**Table S9: primers used in this study.**

| Name | Sequence |
|---|---|
| PfliC-Xho-F | ttactcgagatgcgatttccttttatctttcgacac |
| PfliC-Bam-R | aatggtaccgattcgttatcctatattgcaagtcg |
| PfliC-Z1 | gcatttacgttgacaccatcgaatggcgcaaaacctttcgcggtatgtgtaggctggagctgcttc |
| PfliC-Z2 | ccagtcacgacgttgtaaaacgacggccagtgaatccgtaatcatggtcatgattcgttatcctatattgcaagtc |
| Ptet.flh-P1 | caggtaaatattagtctgattattagtaaagataattaatcaatcactcccgtgtaggctggagctgcttc |
| Ptet.flh-P2 | gtaaatatgacaagttgatgtcataaatgtgtttcagcaactcggaggtatgcatggtacctttctcctctttaatga |
| PfliA-Xho-F | atactcgagacggcaacgccaaattgcctgatg |
| PfliA-Bam-R | aatggatccatagagtgaattcacgataaacagc |
| PfliA-Z1 | gcatttacgttgacaccatcgaatggcgcaaaacctttcgcggtatgtgtaggctggagctgcttc |
| PfliA-Z2 | ccagtcacgacgttgtaaaacgacggccagtgaatccgtaatcatggtcatgataaacagccctgcgttatatgag |

### REFERENCE


1. S. Hui, *et al.*, Quantitative proteomic analysis reveals a simple strategy of global resource allocation in bacteria. *Mol Syst Biol* **11**, e784–e784 (2015).

2. C. You, *et al.*, Coordination of bacterial proteome with metabolism by cyclic AMP signalling. *Nature* **500**, 301–306 (2013).

3. M. Basan, *et al.*, Overflow metabolism in Escherichia coli results from efficient proteome allocation. *Nature* **528**, 99 104 (2015).

4. S. D. Brown, S. Jun, Complete Genome Sequence of Escherichia coli NCM3722. *Genome Announc.* **3** (2015).

5. E. Soupene, *et al.*, Physiological Studies of Escherichia coli Strain MG1655: Growth Defects and Apparent Cross-Regulation of Gene Expression. *J Bacteriol* **185**, 5611–5626 (2003).

6. F. R. Blattner, *et al.*, The Complete Genome Sequence of Escherichia coli K-12. *Science* **277**, 1453–1462 (1997).

7. J. S. Parkinson, Complementation analysis and deletion mapping of Escherichia coli mutants defective in chemotaxis. *Journal of Bacteriology* **135**, 45–53 (1978).

8. C. S. Barker, B. M. Prüß, P. Matsumura, Increased Motility of Escherichia coli by Insertion Sequence Element Integration into the Regulatory Region of the flhD Operon. *Journal of Bacteriology* **186**, 7529–7537 (2004).

9. K. A. Fahrner, H. C. Berg, Mutations That Stimulate flhDC Expression in Escherichia coli K-12. *Journal of Bacteriology* **197**, 3087–3096 (2015).

10. J. Cremer, *et al.*, Chemotaxis as a navigation strategy to boost range expansion. *Nature* **575**, 658–663 (2019).

11. S. Klumpp, Z. Zhang, T. Hwa, Growth Rate-Dependent Global Effects on Gene Expression in Bacteria. *Cell* **139**, 1366–1375 (2009).

12. K. A. Datsenko, B. L. Wanner, One-step inactivation of chromosomal genes in Escherichia coli K-12 using PCR products. *PNAS* **97**, 6640–6645 (2000).

13. G. Kuwajima, Construction of a minimum-size functional flagellin of Escherichia coli. *J Bacteriol* **170**, 3305–3309 (1988).

14. E. Krasnopeeva, C.-J. Lo, T. Pilizota, Single-Cell Bacterial Electrophysiology Reveals Mechanisms of Stress-Induced Damage. *Biophysical Journal* **116**, 2390–2399 (2019).

15. C. Merlin, S. McAteer, M. Masters, Tools for characterization of Escherichia coli genes of unknown function. *Journal of Bacteriology* **184**, 4573–4581 (2002).

16. L. Turner, R. Zhang, N. C. Darnton, H. C. Berg, Visualization of Flagella during Bacterial Swarming. *Journal of Bacteriology* **192**, 3259–3267 (2010).

17. L. Turner, A. S. Stern, H. C. Berg, Growth of Flagellar Filaments of Escherichia coli Is Independent of Filament Length. *Journal of Bacteriology* **194**, 2437–2442 (2012).

18. L. Turner, H. C. Berg, "Labeling Bacterial Flagella with Fluorescent Dyes" in *Bacterial Chemosensing: Methods and Protocols*, Methods in Molecular Biology., M. D. Manson, Ed. (Springer, 2018), pp. 71–76.

19. S. Cayley, M. T. Record, B. A. Lewis, Accumulation of 3-(N-morpholino)propanesulfonate by osmotically stressed Escherichia coli K-12. *Journal of Bacteriology* **171**, 3597–3602 (1989).

20. J. ADLER, B. TEMPLETON, The Effect of Environmental Conditions on the Motility of Escherichia coli. *Microbiology,* **46**, 175–184 (1967).

21. H. C. Berg, L. Turner, Chemotaxis of bacteria in glass capillary arrays. Escherichia coli, motility, microchannel plate, and light scattering. *Biophysical Journal* **58**, 919–930 (1990).

22. J. C. Crocker, D. G. Grier, Methods of Digital Video Microscopy for Colloidal Studies. *Journal of Colloid and Interface Science* **179**, 298–310 (1996).

23. L. Mancini, *et al.*, A General Workflow for Characterization of Nernstian Dyes and Their Effects on Bacterial Physiology. *Biophysical Journal* **118**, 4–14 (2020).

24. J. Rosko, V. A. Martinez, W. C. K. Poon, T. Pilizota, Osmotaxis in Escherichia coli through changes in motor speed. *PNAS* **114**, E7969–E7976 (2017).
